# A Drug-Free Pathogen Capture and Neutralizing Nasal Spray to Prevent Emerging Respiratory Infections

**DOI:** 10.1101/2023.10.02.560602

**Authors:** John Joseph, Helna Mary Baby, Joselyn Rojas Quintero, Devin Kenney, Yohannes A Mebratu, Eshant Bhatia, Purna Shah, Kabir Swain, Shahdeep Kaur, Xiang-Ling Li, John Mwangi, Olivia Snapper, Remya Nair, Eli Agus, Sruthi Ranganathan, Julian Kage, Jingjing Gao, James N Luo, Anthony Yu, Florian Douam, Yohannes Tesfaigzi, Jeffrey M Karp, Nitin Joshi

## Abstract

Respiratory infections pose a global health crisis. Vaccines are pathogen specific, and new vaccines are needed for mutants and emerging pathogens. Here, we report a “drug free” prophylactic platform - a “Pathogen Capture and Neutralizing Spray” (PCANS) that acts *via* a multi-pronged approach to prevent a broad spectrum of respiratory infections. PCANS forms a protective coating in the nasal cavity that enhances the capture of large respiratory droplets. The coating acts as a physical barrier against a broad spectrum of viruses and bacteria, and rapidly neutralizes them, reducing the pathogen load by >99.99%. In mice, PCANS showed nasal retention for at least 8 h and was safe for daily administration. A single prophylactic dose of PCANS protected mice against supra-lethal dosages of a mouse-adapted H1N1 Influenza virus (PR8), reduced lung viral titer by >99.99%, improved survival, and suppressed pathological manifestations. Together, our data suggest PCANS as a promising daily-use prophylactic approach against current and emerging respiratory infections.

## Introduction

Respiratory infections result in significant morbidity and mortality worldwide(*1*). The past few decades have witnessed numerous outbreaks, often leading to epidemics or unanticipated pandemics such as COVID-19. Although vaccines are available against Influenza A virus (IAV), severe acute respiratory syndrome coronavirus 2 (SARS-CoV-2), respiratory syncytial virus (RSV) and *Streptococcus pneumoniae*, the emergence of mutants often reduces the efficacy of vaccines(*2*). Additionally, there are several pathogens, including adenovirus, *Klebsiella pneumoniae, Staphylococcus aureus, and E.Coli,* which can cause severe respiratory diseases, but do not have clinically available vaccines, as of now. In the face of an unforeseen pandemic, the timeline for developing vaccines targeting a pathogen can range from 1 to 10 years, contingent upon the specific nature of the pathogen(*3*, *4*). The rapid creation of efficacious COVID-19 vaccines stands as an unparalleled scientific achievement. However, it took several months for the vaccine to become available, during which numerous hospitalizations and deaths were reported (*5*). Additionally, multiple obstacles, such as production complexities, vaccine nationalism, and the emergence of novel variants, collectively posed major challenges around the world. Another concern pertaining to vaccines is their partial mitigation of the pathogen burden(*6*, *7*), which implies that vaccinated people can still contract and disseminate the infection, albeit at a reduced rate compared to those who are unvaccinated. In addition, a large percentage of the population did not consent to vaccination for various reasons. Thus, there is a critical need to develop a pre-exposure prophylactic approach that can be easily and rapidly employed either independently or in tandem with vaccines, serving as the primary safeguard against current and emerging respiratory pathogens. Such an approach should efficiently reduce pathogen load, and be radically simple to scale up and manufacture to ensure widespread global adoption.

Transmission of most respiratory pathogens predominantly occurs through inhalation of contaminated respiratory droplets and their subsequent deposition in the nasal cavity, which has an entry checkpoint(*8*). For instance, SARS-CoV-2 virus binds to the angiotensin-converting enzyme 2 (ACE2) located in nasal epithelial cells *via* its receptor-binding domain (RBD). The nasal cavity is a primary target for SARS-CoV-2 infection due to high expression of ACE2(*9–11*), which decreases towards the lower respiratory tract(*12*). The infection spreads to the deeper airways *via* virus-laden extracellular vesicles secreted by infected cells in the nasal cavity(*13*). Similarly, bacteria, including *Streptococcus pneumoniae and Staphylococcus aureus* adhere to nasal mucin *via* a specific adhesin receptor(*14*, *15*). Considering the vulnerability of nasal cavity and its critical role in the transmission of respiratory pathogens, chemoprophylactic nasal sprays have been developed to offer pre-exposure prophylaxis against respiratory infections. This approach utilizes chemical agents, including small molecule drugs, antiseptics, or nitric oxide, to deactivate the pathogen in the nasal cavity or a polymer that acts as a physical barrier to prevent pathogen entry through the nasal lining(*16*, *17*). Although multiple pre-exposure chemoprophylactic approaches have been previously developed(*18–20*), they have resulted in suboptimal efficacy with only 20-60% protection achieved in pre-clinical and clinical studies(*21–23*). We contend that the sub-optimal clinical efficacy of previous chemoprophylactic nasal sprays can be attributed, at least in part, to their dependence on a single mode of action, which usually entails either pathogen neutralization or the creation of a physical barrier to hinder pathogen entry through the nasal lining. Furthermore, since pathogens gain access to the nasal cavity through large respiratory droplets(*24*), an optimal prophylactic strategy should also prioritize the effective capture of these droplets laden with pathogens. This necessitates preventing them from bouncing off, a factor that has not been addressed in prior approaches. Previous approaches also consist of a single active ingredient that targets a limited type/class of pathogen(*21*, *25–28*), which could potentially undermine their effectiveness when confronted with newly emerging pathogens. Finally, many chemoprophylactic nasal sprays are unsuitable for repeated/daily application due to toxicity concerns(*29*, *30*).

Herein, we report a Pathogen Capture and Neutralizing Spray (PCANS) that overcomes the aforementioned limitations of previously developed chemoprophylactic nasal sprays, thereby achieving superior efficacy. PCANS has been engineered to act *via* a multi-pronged approach that involves three key steps (Fig. 1). First, PCANS enhances the capture of pathogen-laden respiratory droplets from inspired air by preventing them from bouncing off the nasal lining. To achieve this, we adopted a biomimetic approach that involves reducing the interfacial tension of the nasal lining, similar to pulmonary surfactants in alveoli. Second, PCANS forms a physical barrier over nasal mucosa to intercept invasion/colonization of different pathogens. Last, PCANS consists of multiple “non-drug” agents that rapidly neutralize a wide range of pathogens. We define ‘neutralization’ as a process that impedes pathogen entry into host cells by either destabilizing the pathogen cell membrane or blocking the receptor-mediated binding/fusion of the pathogen through chemical interactions. To ensure safety during daily or repeated use, PCANS was meticulously designed as a “drug-free” formulation, incorporating biopolymers, surfactants, and alcohols that are listed in the inactive ingredient database (IID) or generally recognized as safe (GRAS) list of the Food and Drug Administration (FDA), and are present as excipients in commercially available nasal/topical formulations. These components and their unique concentrations were identified *via* a highly iterative approach aimed at maximizing sprayability, mucoadhesiveness, the capture of respiratory droplets, physical barrier property, pathogen neutralization activity, and nasal residence time. *In vitro*, PCANS demonstrated excellent physical barrier properties against multiple viruses and bacteria, and rapidly neutralized them, resulting in >99.99% reduction in the pathogen load. Coating a 3D-model nasal cavity with PCANS significantly increased the capture of large respiratory droplets, compared to only a mucus-coated nasal cavity. Intranasal administration of PCANS-loaded with a fluorescent dye resulted in at least 8 h of residence time in the mouse nasal cavity, measured as the retention of fluorescence signal over time. In a proof-of-concept *in vivo* study performed in a mouse model of severe Influenza A infection induced by a supra-lethal dose of PR8 virus (a mouse-adapted H1N1 Influenza virus), we demonstrated that pre-exposure prophylactic administration of a single dose of PCANS resulted in >99.99% reduction in lung viral titer and 100% survival as compared to 0% observed in the PBS-treated group. Overall, PCANS holds potential as a promising pre-exposure prophylactic approach to prevent current and emerging respiratory infections. The uncomplicated and easily scalable manufacturing process of PCANS, coupled with its “drug free” nature and robust stability, as demonstrated in this study, renders it conducive for global distribution and widespread adoption.

**Fig. 1:**
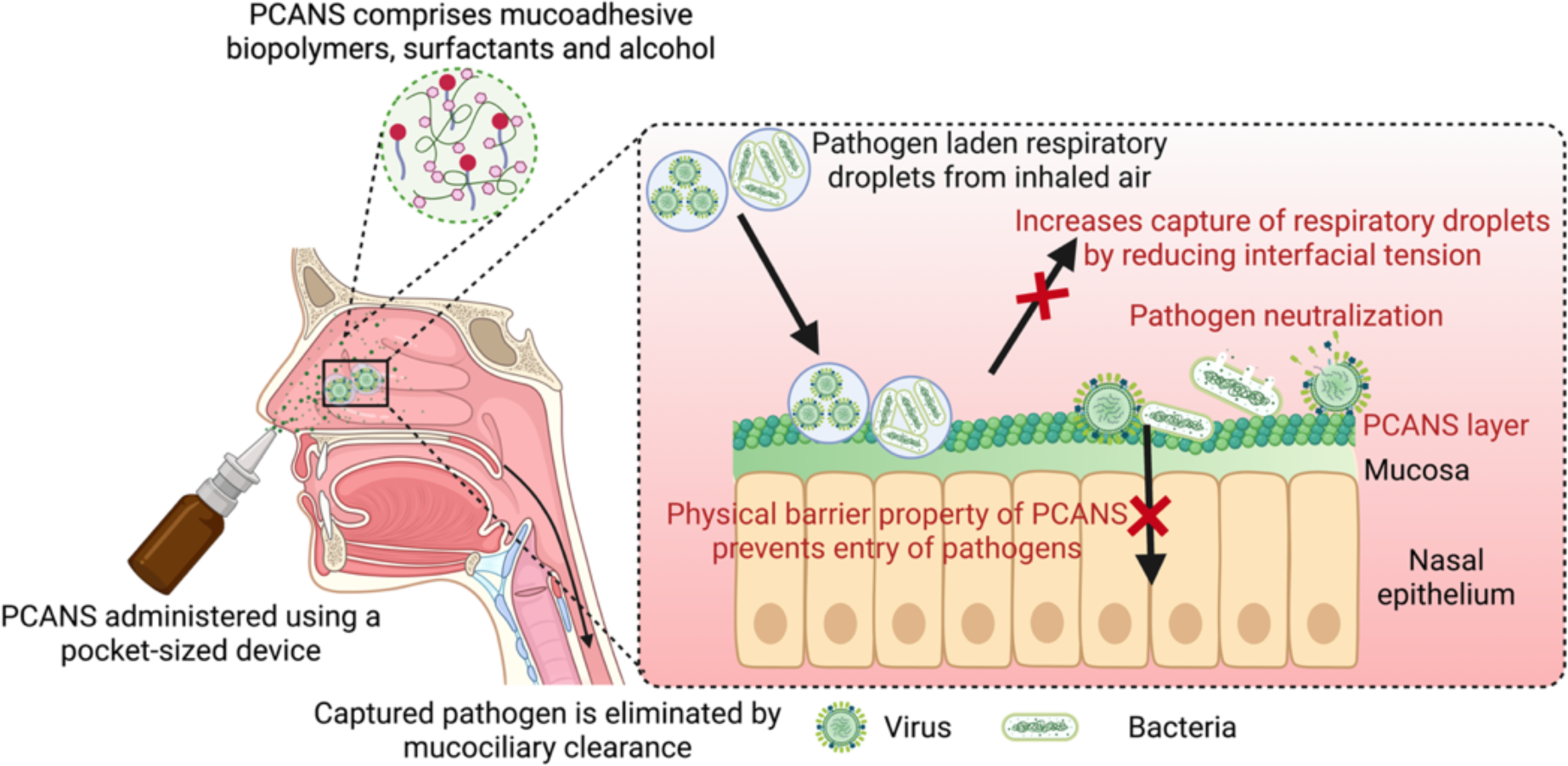
Pathogen Capture and Neutralizing Spray (PCANS) acts *via* a multi-pronged approach against respiratory pathogens. An aqueous, “drug-free” solution of PCANS, comprising mucoadhesive biopolymers, surfactants, and alcohol, is administered using a pocket-sized nasal spray device and undergoes a phase transition to form a hydrogel layer over nasal mucosa. Surfactants in PCANS reduce interfacial tension of the nasal lining and increase wettability to enhance the capture or reduce the bounce-off of pathogen-laden respiratory droplets from the inhaled air. PCANS layers as a physical barrier preventing the transport of pathogens through the nasal lining. Finally, pathogens are neutralized by biopolymers and surfactants present in PCANS. PCANS is cleared *via* the native mucosal clearance mechanism and is eliminated through the digestive route.

## Results

### Leveraging biopolymers to restrict pathogen entry *via* formation of a physical barrier

We selected mucoadhesive biopolymers that are listed in the IID or GRAS list of the FDA and are present as excipients in commercially available nasal/topical formulations. Specifically, gellan, pectin, hydroxypropyl methylcellulose (HPMC), carboxymethyl cellulose sodium salt (CMC), carbopol, and xanthan gum were selected. The biopolymers were screened for their ability to impart physical barrier property to PCANS. Since a metered spray device would be used to administer PCANS, we first identified sprayable concentration of each biopolymer by performing rheological measurements (Fig. 2 a-f). Dynamic viscosity curves were generated using a rotational rheometer by varying shear rates up to 40 s^−1^, which is within the lower limits of shear rates encountered while dispensing formulations through a nasal spray device. Concentrations that exhibited a viscosity of less than 0.1 Pa.s were considered ‘sprayable’ (*31*, *32*). Next, we determined the mechanical strength of each biopolymer at the highest sprayable concentration before and after the addition of simulated nasal fluid (SNF). SNF was added to mimic the physiological environment in the nasal cavity. Mechanical strength was measured using a rotational rheometer and quantified as storage modulus (G’), which represents the amount of structure present in a material(*33*). In the presence of SNF, gellan showed the highest G’ as compared to other biopolymers (Fig. 2g), indicating its superior mechanical strength. Gellan showed a 100-fold increase in its G’ in the presence of SNF (Fig. 2g), which is consistent with its ability to undergo *in situ* gelation under physiological conditions. Mono and divalent cations present in the SNF complex with glucuronic monomeric units of gellan to form a crosslinked hydrogel(*34*). Compared to gellan, other biopolymers showed minimal or no increase in their storage modulus, suggesting poor *in situ* gelation. To investigate physical barrier property of biopolymers, a trans-membrane assay was devised (Fig. S1), which involved evaluating the transport of IAV through an SNF-coated cell strainer (pore size ∼70 µm) or a cell strainer coated with simulated mucus/SNF mixture or a biopolymer/SNF mixture. After 4 h, the viral titer in the chamber below the strainer was quantified by performing a plaque assay in Madin-Darby canine kidney (MDCK) host cells. Consistent with its excellent mechanical strength, Gellan/SNF reduced the transport of IAV particles by >4-log fold (99.99%) as compared to only SNF-coated or mucus/SNF-coated strainers (Fig. 2h). Xanthan/SNF, CMC/SNF and HPMC/SNF also significantly reduced the IAV transport, but not as efficiently as gellan/SNF. Interestingly, despite significantly lower mechanical strength of pectin/SNF as compared to gellan/SNF, it intercepted the IAV transport with similar efficiency as gellan/SNF. Carrageenan, a biopolymer used in previously reported and commercially available chemoprophylactic nasal sprays(*35*), was used as a control and did not reduce IAV transport in the presence of SNF.

**Fig. 2:**
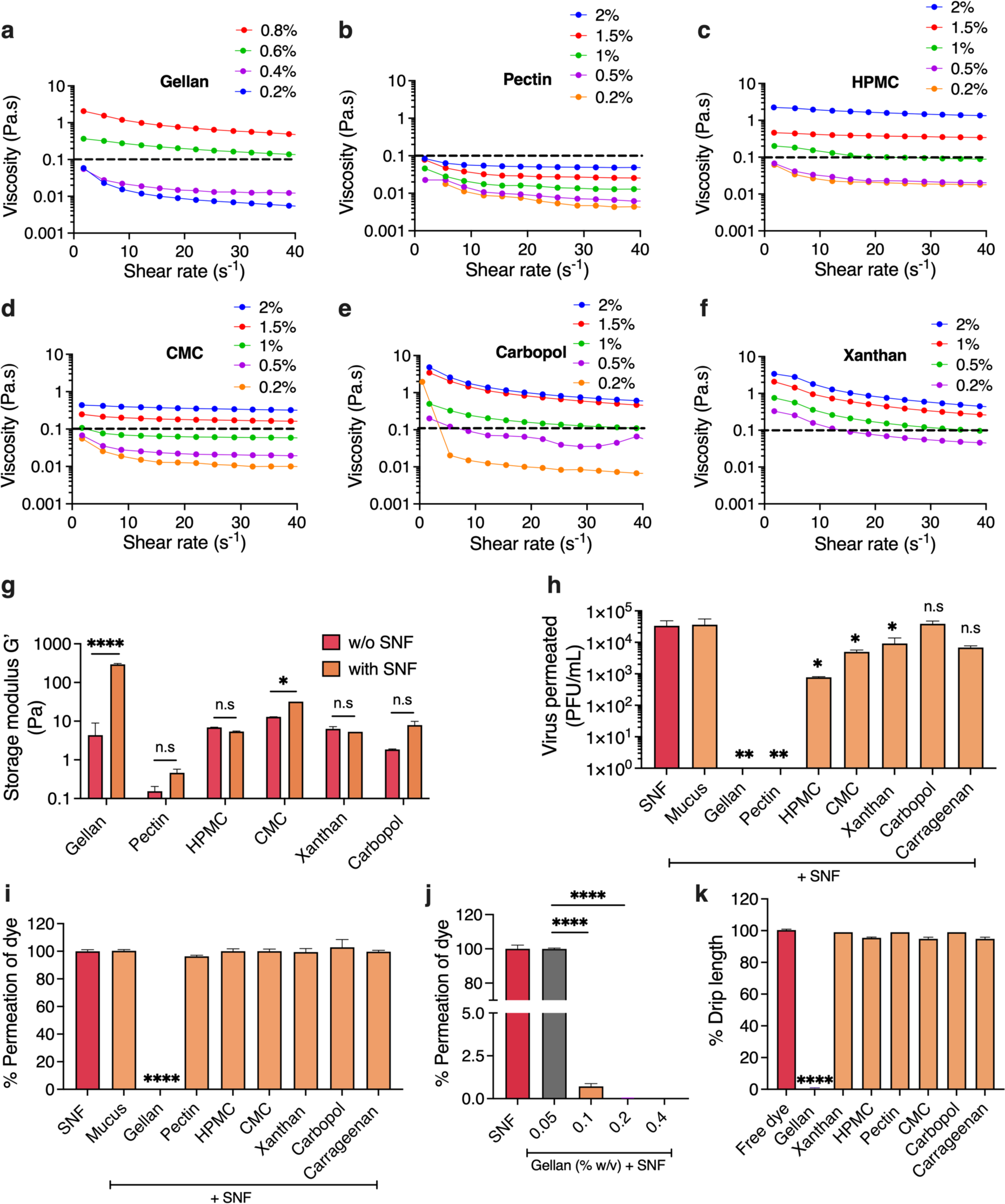
Biopolymers were screened for physical barrier property against pathogen entry. Viscosity as a function of shear rate up to 40 s^−1^ at 25°C for different concentrations of (**a)** gellan, **(b)** pectin, **(c)** hydroxy propyl methyl cellulose (HPMC), **(d)** carboxymethylcellulose (CMC), **(e)** Carbopol and **(f)** xanthan gum in water. The sprayable viscosity window is shown below the dashed line. **(g)** Storage modulus (G’) of 0.4% (w/v) gellan, 2% (w/v) pectin, 0.5% (w/v) HPMC, 0.5% (w/v) CMC, 0.2% (w/v) Carbopol, and 0.2% (w/v) xanthan gum, without and with simulated nasal fluid (SNF). Amplitude sweep measurements were performed at 37°C by varying oscillatory strain between 0.005% to 10 % at 1 Hz frequency. \*\*\*\**P* < 0.0001, \**P* < 0.05. n.s., not significant. **(h)** Amount of Influenza A virus (IAV) that permeated within 4 h through a simulated nasal fluid (SNF)-coated cell strainer (pore size ∼70 µm) or a cell strainer coated with simulated mucus/SNF mixture or a biopolymer/SNF mixture. Permeation of viral particles was quantified by evaluating the viral titer in the chamber below the strainer using plaque assay performed in MDCK host cells. Results are expressed in plaque-forming units (PFU/mL). \*\**P* < 0.01, \**P* < 0.05 compared to mucus/SNF, n.s, not significant. Percentage permeation of a fluorescent dye, rhodamine B isothiocyanate through **(i)** an SNF-coated cell strainer or a cell strainer coated with simulated mucus/SNF mixture or a biopolymer/SNF mixture. \*\*\*\**P* < 0.0001 compared to mucus/SNF and **(j)** an SNF-coated strainer or strainer coated with gellan/SNF at different concentrations of gellan. \*\*\*\**P* < 0.0001 compared to 0.05% w/v gellan/SNF. **(k)** Percentage drip length of free brilliant green dye or mucoadhesive polymers mixed with brilliant green dye on porcine mucosal tissue. Drip length from the spray area was measured as the distance traversed in 4 h by the biopolymer or free dye from the point of deposition. The percentage drip length of each biopolymer was calculated with respect to the drip length of the free dye. \*\*\*\**P* < 0.0001 compared to free dye. For **g and h**, *P*-values were determined using two-way ANOVA with Tukey’s multiple comparisons tests. For **i-k**, *P*-values were determined using one-way ANOVA with Tukey’s post hoc analysis. Data in **a-f** are from a single experiment (experiment repeated three times). Data in **g-k** are means ± SD of technical repeats (n = 3, each experiment performed at least twice).

Reduction in the transport of IAV particles by anionic biopolymers could be a result of their physical barrier property and/or electrostatic interactions between their negatively charged polymeric chains and the positively charged capsid of IAV. To decouple the effects of physical barrier property and electrostatic interactions, we studied the transport of a low molecular weight anionic dye, rhodamine B isothiocyanate, that would abate electrostatic interactions with the anionic biopolymers. Gellan/SNF resulted in 100% reduction in the transport of the dye, confirming excellent physical barrier property (Fig. 2i). Other biopolymers did not reduce the transport of the dye, indicating their poor physical barrier property. This also indicates that the reduction in IAV transport by pectin was primarily mediated *via* electrostatic interaction of pectin’s chains with the virus capsid. Interestingly, gellan/SNF reduced the transport of rhodamine B dye in a concentration-dependent manner (Fig. 2j), which was consistent with the concentration-dependent increase in the mechanical strength (G’) of gellan in the presence of SNF (Fig. S2). A 0.2% w/v concentration of gellan also reduced the transport of *E.coli* bacteria by >8-log fold (100%) (Fig. S3), suggesting it’s broad spectrum physical barrier property to limit the transport of both viruses and bacteria. Mucus, on the other hand, only showed a 1-log fold (90%) reduction. To conclude, gellan at a concentration of 0.2% w/v and above impeded the transport of rhodamine B dye, *E. coli*, and IAV by 100%.

To ensure maximum coverage of the nasal cavity, we evaluated the spray characteristics of gellan at a concentration of 0.2% w/v or higher with a hydraulic spray nozzle. Plume geometry and spray coverage were measured with a high-speed image acquisition system. Increasing gellan concentration resulted in a significant reduction in the angle of emitted plume of the spray (defined as ‘plume angle’) and coverage area (Fig. S4a-d).

Next, we evaluated the retention ability of gellan and other biopolymers at the mucosal tissue upon spraying. Mucosal retention was measured as the drip length, defined as the distance traversed in 4 h by the biopolymer from the point of deposition on sheep’s intestinal mucosa placed vertically. To visualize dripping, biopolymers were mixed with a brilliant green dye. The percentage drip length of each biopolymer was calculated with respect to the drip length of the free dye. Gellan demonstrated excellent mucosal retention with zero drip length (Fig. 2k and Fig. S5). Other biopolymers, including carrageenan, which was used as a control showed >95% drip length, indicating poor mucosal retention. Gellan’s superior mucosal retention is attributed to its ability to strongly entangle with mucin glycoprotein in the mucosal tissue during the sol-gel transition(*36*).

### Identifying agents for neutralizing a broad-spectrum of respiratory pathogens

To impart PCANS a broad-spectrum pathogen neutralization ability, we screened agents from three different classes of compounds, including biopolymers, surfactants, and alcohols. These compounds were selected based on their previously reported ability to neutralize different types of pathogens(*37–39*). To maximize safety and translatability of PCANS, we only selected agents that are listed in the IID or GRAS list of the FDA and are present as excipients in commercially available nasal/topical formulations (Fig. 3a). We first evaluated the neutralization ability of these agents against viruses. Neutralization was studied *in vitro* by incubating each agent individually with either IAV or SARS-CoV-2 for 10 or 60 min, followed by 1-min centrifugation and subsequent infection of target cells with the supernatant evaluated using plaque forming or focus-forming assay. We chose IAV and SARS-CoV-2 due to their high prevalence worldwide as respiratory viruses and also due to a difference in their capsid proteins and charge(*40*, *41*). Biopolymers were evaluated at their highest sprayable concentration, except for gellan and carrageenan. Gellan was evaluated at 0.2% w/v due to its superior physical barrier property compared to 0.1% w/v concentration and superior spray pattern compared to 0.4% w/v concentration. Carrageenan, used as a control, was evaluated at 0.16% w/v, as this concentration is present in a commercially available chemoprophylactic nasal spray(*42*, *43*). Surfactants and alcohols were evaluated at the highest concentration previously used in humans *via* nasal route(*44*, *45*). Compared to carrageenan, pectin exhibited superior neutralization of IAV, regardless of the incubation time, and demonstrated a 4-log fold (99.99%) reduction in viral titer in the host cells in comparison to PBS (Fig. 3b). Ten min of incubation with carbopol did not reduce the IAV titer, but a 4-log fold (99.99%) reduction was observed with 60 min of incubation. Gellan exhibited similar neutralization of IAV as carrageenan, resulting in only a 1-log fold (90%) reduction in viral load in the host cells. For SARS-CoV-2, both pectin and carrageenan showed less than a 1-log fold decrease in viral load in the host cells (Fig. 3g). Gellan showed a 4-log fold (99.99%) reduction in the viral titer, but only with 1 h incubation time. Among surfactants, tween 80 and benzalkonium chloride (BKC) showed a 1-log log fold reduction in IAV titer in the host cells, regardless of the incubation time (Fig. 3c). Rapid neutralization of SARS-CoV-2 was observed with BKC, resulting in a 5-log fold (>99.99%) reduction in viral load in the host (Fig. 3h). Alcohols did not neutralize SARS-CoV-2, and minimum neutralization was observed for IAV, resulting in less than 1-log fold (90%) reduction in viral load for chlorobutanol and phenethyl alcohol (PEA) (Fig. 3d and 3i). Overall, this extensive screening identified pectin and BKC as the most effective agents for rapid neutralization of IAV and SARS-CoV-2, respectively. Neutralization ability of pectin and BKC was found to be dose-dependent (Fig. 3e and 3j).

**Fig. 3:**
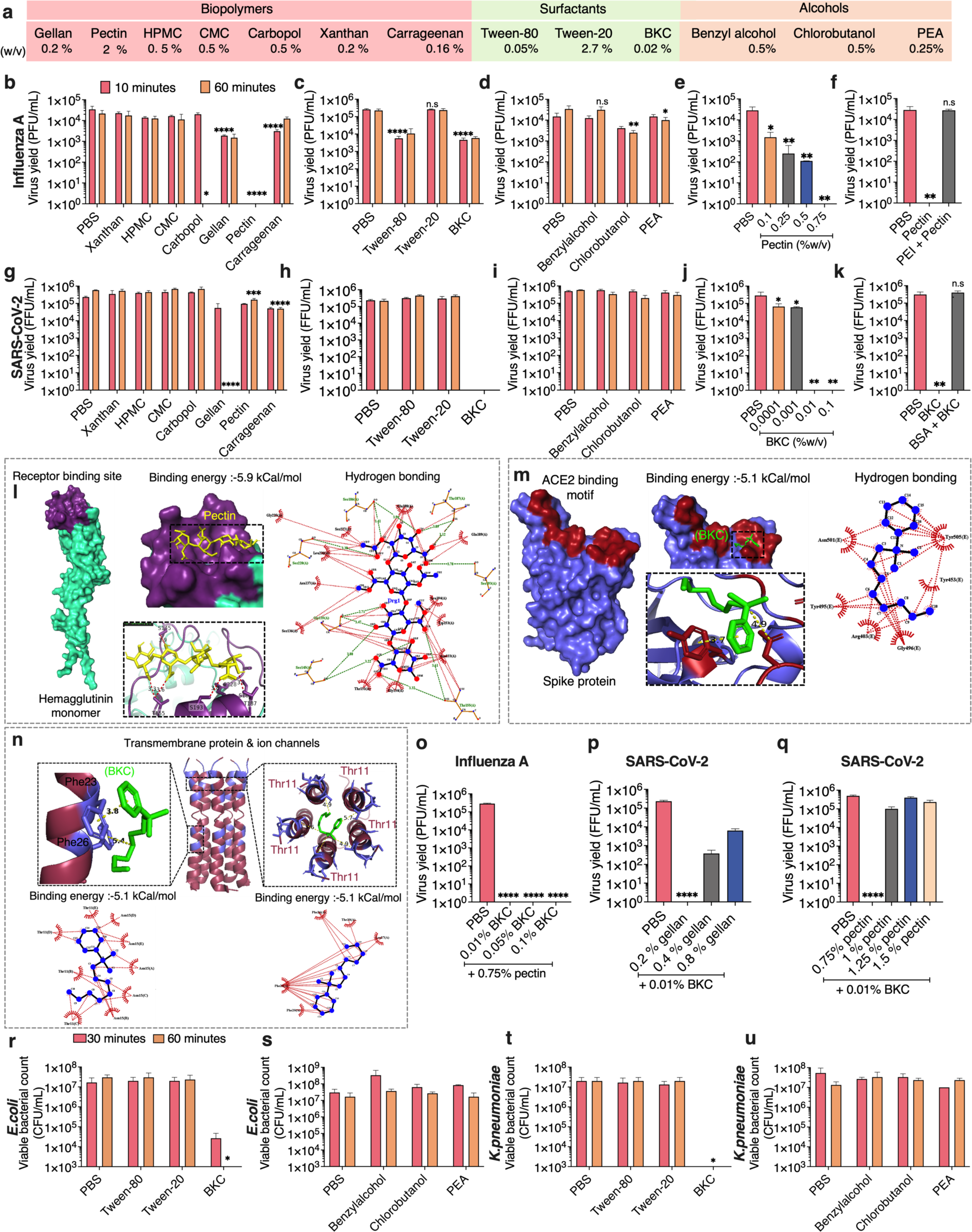
Biopolymers, surfactants and alcohols were screened for neutralization of different respiratory pathogens. **(a)** Table summarizes different components and their concentrations to determine the neutralization ability against respiratory pathogens. Each component was individually evaluated for its pathogen neutralization potential. IAV and SARS-CoV-2 viral loads in the host cells after 10 or 60 min incubation of the virus with **(b, g)** different biopolymers, **(c, h)** different surfactants, and **(d, i)** different alcohols. Viable viral titer was quantified using plaque assay in MDCK host cells for IAV and focus-forming assay in Vero E6 cells for SARS-CoV-2 virus. Results are expressed in plaque-forming units (PFU/mL) or focus-forming units (FFU/mL). \*\*\*\**P* < 0.0001, \*\*\**P* < 0.001, \*\**P* < 0.01, \**P* < 0.05 compared to 10 minutes of incubation with PBS. n.s, not significant. Viral loads in the host cells after 10 min incubation of **(e)** IAV and **(j)** SARS-CoV-2 with different concentrations of pectin and BKC, respectively. \*\**P* < 0.01, \**P* < 0.05 compared to PBS. Viral loads in the host cells after 10 min incubation of **(f)** IAV and **(k)** SARS-CoV-2 with pectin + polyethylenimine and BKC + bovine serum albumin, respectively. \*\**P*<0.01 compared to PBS. n.s, not significant. **(l)** Pectin (yellow) binds to the receptor binding site of IAV (purple) at the distal part of hemagglutinin monomer (colored in purple) through hydrophobic interactions with Ser227 and Glu190, and hydrogen bonding with Ser228, Ser186, and Thr187. Blue and red dots in hydrogen bonding maps represent carbon and oxygen atoms, respectively. **(m)** Chemical interaction of BKC (green) with ACE2 binding motif (red) in the spike protein of SARS-CoV-2. Interaction map reveals the hydrogen bonding of BKC with Tyr505 and Gly496. **(n)** Aromatic pi-pi interaction of BKC (green) with Phe23 (purple) in the transmembrane domain and with Thr11 (brown) membrane helices. Interaction analysis shows 10 hydrophobic bonds with Phe23 and 8 hydrophobic bonds with Phe26. **(o)** Viral load in host cells after 10 min incubation of IAV with pectin in the presence of different concentrations of BKC. \*\*\*\**P* < 0.0001 compared to PBS. Viral load in the host cells after 10 min incubation of SARS-CoV-2 with BKC (0.01% w/v) in the presence of different concentrations of **(p)** gellan and **(q)** pectin. \*\*\*\**P* < 0.0001 compared to PBS. Effect of surfactants **(r, t)** and alcohols **(s, u)** against gram-negative bacteria *E. coli* and *K. pneumoniae* using colony-forming unit (CFU) plate count method after 30 and 60 minutes of exposure. Viable bacterial colonies are expressed in CFU/mL. \**P*<0.05 compared to PBS. For **b-d**, *P* values were determined using two-way ANOVA with Tukey’s post hoc analysis. For **e, f, j, k, o-q**, *P* values were determined using one-way ANOVA. Data in **b-k** and **o-u** are presented as Means ± S.D of technical repeats (n = 3, each experiment performed at least twice). Ser, serine; Thr, threonine; Glu, glutamic acid; Phe, phenylalanine; Tyr, tyrosine; Gly, glycine.

To elucidate the viral neutralization mechanism of pectin and BKC, we performed *in silico* modeling to determine their binding affinity with the receptor binding domains (RBD) of IAV and SARS-CoV-2, respectively. For IAV, anionic pectin targets RBD at the distal part of hemagglutinin, which is positively charged, thus averting the virus entry into the host cell (Fig. 3l). Compared to the host ligand sialic acid present in mucin, pectin showed stronger binding to RBD through distant hydrogen bonding with Se228, Ser186, and Thr187 and hydrophobic linkage with Ser227 and Glu190 (Fig. S6). BKC was found to exhibit hydrophobic interactions with the ACE2 binding motif of spike protein of SARS-CoV-2 (Fig. 3m). BKC also showed hydrophobic interactions with Phe23 and Phe26 in membrane helices via *pi*-*pi* stacking (Fig. 3m), which can distort the helical conformation of adjacent helices, as aromatic stacking of Phe23 and Phe26 is a prerequisite to stabilizing helix-helix interface of the envelope transmembrane protein. BKC fits into the pentameric ion channels at the N terminus of the transmembrane domain through interaction with Thr11 and potentially blocks the influx/efflux of ions (Fig. 3n). To determine the role of electrostatic interaction in pectin- and BKC-mediated neutralization of IAV and SARS-CoV-2, respectively, we performed a neutralization assay by pre-treating pectin and BKC with counter ions to offset the charge. As anticipated, anionic pectin in the presence of positively charged polyethyleneimine lost its neutralization activity and failed to show a significant reduction in the viral load compared to PBS (Fig. 3f). Likewise, the pretreatment of BKC with negatively charged bovine serum albumin diminished the ability of BKC to reduce the SARS-CoV-2 titer in the host cells (Fig. 3k).

Next, we investigated whether ionic interactions between anionic gellan or pectin with cationic BKC, when present together in a formulation, would impact the neutralization ability of pectin or BKC. Notably, the neutralization efficiency of pectin (0.75% w/v) against IAV remained conserved even with a dose-dependent increase in BKC up to a concentration of 0.1% w/v (Fig. 3o). Neutralization efficiency of BKC (0.01% w/v) against SARS-CoV-2 was not impacted by gellan or pectin at 0.2% w/v or 0.75% w/v concentrations, respectively, but reduced at higher concentrations (Fig. 3p,q). These results further underscore that the concentration of each agent is critical for efficient neutralization.

Finally, we also screened surfactants and alcohols to assess their neutralization ability against bacteria, including *E. coli* and *Klebsiella pneumoniae (K. pneumoniae)*. Neutralization was determined by measuring the bactericidal activity. Each agent was individually incubated with either *E. coli* or *K. pneumoniae* for 30 or 60 min, followed by 1-min centrifugation, and then evaluating the bacterial load in the supernatant using a colony-forming assay. BKC was more effective than non-ionic surfactants, resulting in a 4-log fold (99.99%) and 7-log fold (99.99%) reduction in colony-forming units (CFU) of *E. coli* and *Klebsiella pneumoniae*, respectively, with an incubation time of 30 min (Fig. 3r, t). Alcohols had a negligible bactericidal effect over the exposure periods of 30 or 60 min (Fig. 3s,u). Altogether, our data on physical barrier property, spray pattern, mucosal retention, and neutralization indicate gellan, pectin, and BKC as the three critical components to formulate PCANS. However, we also incorporated phenethyl alcohol (PEA), as it is commonly added as a stabilizer to nasal formulations to prevent the growth of gram-negative bacteria and fungi (*44*) and ensure long shelf life.

### Utilizing surfactants to promote the capture of respiratory droplets

Pulmonary surfactant layers the alveolar epithelium to enhance wettability and trap airborne particles(*46*). We adopted this biomimetic approach to capture pathogen-laden respiratory droplets in the nasal cavity. Specifically, we identified surfactants to reduce interfacial tension of PCANS and reduce the bounce off/escape of respiratory droplets. We evaluated surfactants listed in the IID list, including Tween-20, Tween-80, and BKC. Screening was performed using a twin impinger, which is a glass apparatus that can be used to assess the deposition of aerosolized particles in different regions of the respiratory tract (*47*, *48*)(Fig. 4a). Simulated mucus or a biopolymer mixture of gellan and pectin without or with different concentrations of surfactants was sprayed into the SNF-coated oropharyngeal region of the impinger (Fig. 4a). Droplets with mass medial aerodynamic diameter >5 µm and laden with rhodamine B-loaded liposomes (size ∼400 nm) were generated using a jet nebulizer to mimic pathogen-laden large respiratory droplets. Droplet capture was determined by quantifying the fluorescence intensity of rhodamine B in the biopolymer/surfactant mixture or the mucus layer. Biopolymer mixture without any surfactant showed similar fluorescence intensity as mucus (Fig. 4b). Combining the biopolymer mixture with Tween-80 or Tween-20 at a concentration higher than 0.005% w/v or with BKC at a concentration higher than 0.01% w/v resulted in a significant increase in the fluorescence intensity as compared to mucus or only biopolymer mixture, suggesting increased capture of droplets due to surfactants. Compared to Tween-20, BKC and Tween-80 resulted in a significantly higher fold increase in the fluorescence intensity when added to the biopolymer mixture at a concentration of 0.05% w/v or higher (Fig. 4b). At 0.05% w/v concentration, both BKC and Tween-80 containing biopolymer mixtures showed similar fluorescence intensity, which was 4-fold higher than the fluorescence intensity of mucus or biopolymer mixture without a surfactant. Since 0.01 % w/v is the most commonly used concentration of BKC in commercially available nasal formulations(*49*, *50*), and also showed excellent neutralization activity against SARS-CoV-2, we decided to use this concentration, even though BKC didn’t increase the capture of respiratory droplets at this concentration. To impart respiratory droplet-capturing ability, we decided to proceed with Tween-80 and determined its safe concentration that would not compromise the permeability or metabolic activity of nasal epithelium. To that end, we performed an *in vitro* assay evaluating the transepithelial electrical resistance (TEER) across the human nasal epithelial cell (RPMI-2650)-based monolayer upon treatment with different concentrations of tween-80. A transient dip of less than 15% in TEER was observed in the monolayer immediately after the addition of Tween-80, irrespective of the concentrations evaluated in this study. However, the TEER reversed rapidly to the original value in less than 1 h after replacing tween-80-containing medium with fresh medium (Fig. 4c). The drop in the TEER for Tween-80 was significantly less compared to Triton-X (negative control), which resulted in a permanent change in the TEER. Second, we evaluated the effect of different concentrations of Tween-80 on the metabolic activity of RPMI-2650 cells upon 24 or 48 h of incubation. Cells incubated with 0.01% or 0.05% w/v tween-80 showed similar metabolic activity as cells incubated in medium. However, Tween-80 (0.5% w/v) resulted in a significant reduction in the metabolic activity of RPMI cells (Fig. S7). Overall, based on our data for physical barrier property, spray pattern, mucosal retention, neutralization, droplet capture, and nasal epithelial cell toxicity, we decided on gellan, pectin, BKC, PEA, and Tween-80 as the final components for PCANS, and validated the respiratory droplet capturing ability of the final formulation using a 3D-model of human nasal cavity (Koken cast) with the anatomical intricacies (*51*) (Fig. 4d). Consistent with the twin impinger results, there was no significant difference in the fluorescence intensity between gellan and pectin mixture, and mucus (Fig. 4e). PCANS, on the other hand, showed a 2-fold higher fluorescence compared to mucus, suggesting the potential of PCANS to increase the capture of pathogen-laden respiratory droplets from inhaled air.

**Fig. 4:**
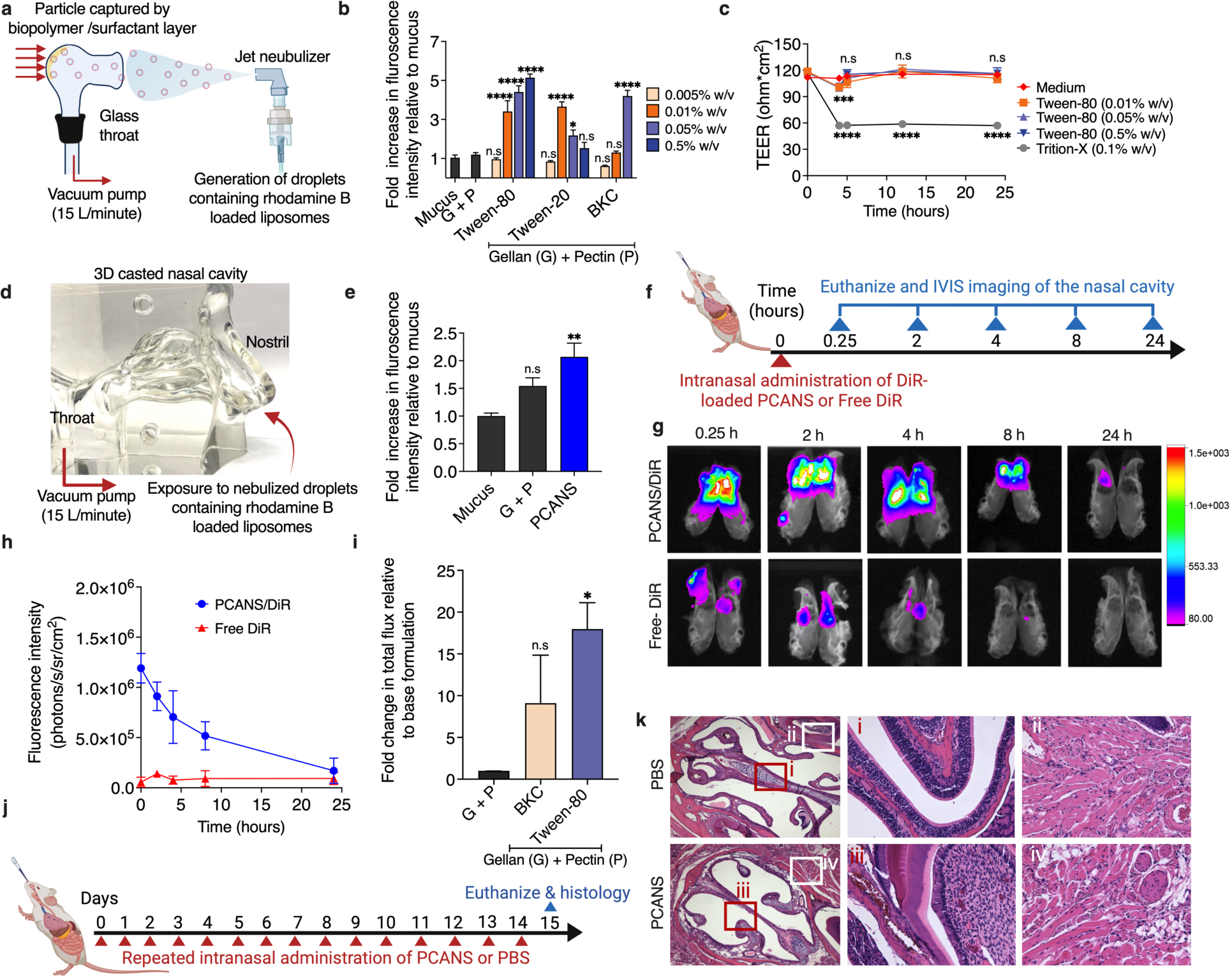
PCANS enhances the capture of respiratory droplet-mimicking aerosol and exhibits prolonged nasal residence time in mice. **(a)** Experimental design for measuring the capture of respiratory droplet-mimicking aerosol. A twin impinger was used to simulate the aerodynamics of the human respiratory tract. Mucus or gellan and pectin solution (G+P), without or with different concentrations of Tween-80, Tween-20 or BKC was coated on the inner surface of the throat region of the impinger using a nasal spray device. Droplets with mass medial aerodynamic diameter >5 µm and laden with rhodamine B-loaded liposomes (size ∼400 nm) were generated using a jet nebulizer and administered into the impinger under vacuum (15 L/min). Droplet capture was determined by quantifying the fluorescence intensity of rhodamine B in the biopolymer/surfactant mixture or mucus layer. (**b)** Fold increase in fluorescence intensity with respect to mucus. \*\*\*\**P* < 0.0001, \**P* < 0.05 compared to mucus. (**c)** Transepithelial electrical resistance (TEER) across the human nasal epithelial cell (RPMI-2650)-based monolayer at different time points after treatment with only medium or medium containing Triton-X (0.1% w/v) or different concentrations of Tween-80. Surfactant-containing medium was replaced at 4 h with fresh medium to examine impedance recovery. \*\*\*\**P* < 0.0001, \*\*\**P* < 0.001 compared to untreated control. n.s, not significant. (**d)** Experimental design for measuring the capture of respiratory droplet-mimicking aerosol using a 3D human nasal cavity model (Koken cast). The inner surface of the nasal cavity was coated with mucus, G+P solution or PCANS (the final formulation) using a nasal spray device. The throat part of the model was coupled to a vacuum pump for simulating the respiratory airflow (15 L/min). Nostrils were then exposed to nebulized rhodamine B-loaded liposomes for 1 minute. Droplet capture was determined by quantifying the fluorescence intensity of rhodamine B in the nasal cavity. **(e)** Fold increase in fluorescence intensity with respect to mucus. \*\**P* < 0.01 compared to mucus. n.s, not significant. **(f)** Experimental outline for the evaluation of the nasal residence time of PCANS in mice. C57Bl/6 mice were intranasally administered with 10μL of free DiR or DiR-loaded PCANS (PCANS/DiR) into each nostril. Mice were euthanized at different time points over 24 h, and nasal cavity was harvested and imaged using an *in vivo* imaging system (IVIS). (**g)** Representative images of the nasal cavity excised at different time points. (**h)** Quantification of fluorescence intensity in the nasal cavity at different time points. (**i)** Fold change in total flux at 8h in the nasal cavity relative to G+P. \**P* < 0.05, compared to G+P. n.s, not significant. (**i)** Experimental design to assess the biocompatibility of PCANS in mouse nasal cavity. 10 µl PCANS or PBS was administered into each nostril of C57Bl/6 mice once daily for 14 consecutive days. Animals were euthanized on day 15 and nasal cavity was analyzed histologically. (**j)** Representative images of H&E-stained sections of nasal turbinate from mice captured using a 4X objective. Insets represent healthy olfactory epithelium (i) and (iii), and lamina propria (ii) and (iv) captured at 20X objective. For **b, e and i***, P* values were determined by one-way ANOVA using Tukey’s post hoc analysis. For **b**, concentrations for each surfactant were compared individually. For **c**, *P* values were determined by two-way ANOVA with Tukey’s multiple comparison test. Data in **b,c,** and **e** are presented as Means ± S.D of biological repeats (n = 3, each experiment performed at least twice). Data in **h** and **i** are presented as Means ± SEM (n=5 mice/group).

### Prolonged nasal retention of PCANS and safety upon repeated administration

Next, we evaluated the retention of PCANS in the nasal cavity of mice (Fig. 4f). PCANS (10 µL) mixed with a fluorescent dye – (DiIC18(7) (1,1’-Dioctadecyl-3,3,3’,3’-Tetramethylindotricarbocyanine Iodide) (DiR) was administered into both nostrils of C57/BL6 mice. Free DiR was used as a control. Mice were euthanized at different time points over 24 h, and nasal cavity was harvested and imaged using an *in vivo* imaging system (IVIS) to quantify the fluorescence signal from DiR. Free DiR resulted in negligible fluorescence signal, even at 15 min after administration, suggesting its rapid clearance (Fig. 4g,h). Interestingly, mice administered with DiR-loaded PCANS showed significant fluorescence for up to 8 h, suggesting prolonged nasal retention of PCANS (Fig. 4g,h). We hypothesized that prolonged retention of PCANS is attributed to the presence of surfactants, including Tween-80 and BKC, which have previously been shown to reduce cilia beat frequency in the nasal cavity(*52*). To test our hypothesis, we compared nasal retention of DiR-loaded mixture of gellan and pectin without or with tween-80 or BKC. The addition of both BKC or tween-80 significantly enhanced the nasal retention of gellan and pectin mixture at 8 h post-nasal administration, as evident from the fluorescent signal of DiR in the nasal cavity (Fig. 4i). However, tween-80 resulted in significantly higher nasal retention than BKC. The ability of tween-80 to enhance the nasal retention of gellan and pectin mixture was found to be concentration-dependent (Fig. S8). However, considering irreversible nasal epithelial permeabilization and cytotoxicity at 0.5% w/v or higher concentration of tween-80, we maintained 0.05% w/v in PCANS for further experiments. Notably, nasal administration of DiR-loaded PCANS only showed fluorescence signals in the nasal cavity and stomach, suggesting no systemic absorption. PCANS was fully cleared at 24 h (Fig. 4g,h, and S9), resulting in negligible fluorescence signal in both the nasal cavity and the stomach. To confirm safety of PCANS, we performed a repeat-dose toxicity study in healthy mice intranasally administered with PCANS or PBS once daily for 14 consecutive days (Fig. 4j). Hematoxylin and eosin (H&E) stained sections of nasal cavity from both PBS or PCANS-administered mice did not show any inflammation or other gross evidence of toxicity, as evident by a defined lamina propria (Fig. 4k). This connotes the safety of PCANS for daily administration.

### Broad-spectrum activity, spray characteristics, and shelf stability of PCANS

Having identified the final components of PCANS, along with their optimal concentrations, we sought to demonstrate the physical barrier property and neutralization ability of PCANS against a broad spectrum of respiratory pathogens, including enveloped viruses (IAV, SARS-CoV-2, RSV), a non-enveloped virus (adenovirus), and bacteria (*E. Coli* and *K. Pneumonaie*). Physical barrier property was evaluated by assessing the transport of pathogens through an SNF-coated cell strainer or a cell strainer coated with simulated mucus/SNF mixture or PCANS/SNF mixture. PCANS/SNF prevented the transport of all the pathogens by >4-log fold (>99.99%) (Fig. 5a-f), suggesting its broad-spectrum physical barrier property. For all pathogens, except RSV, mucus/SNF mixture showed significantly less prevention of pathogen transport compared to PCANS/SNF. PCANS also efficiently neutralized all the tested pathogens within 10 min of incubation time, resulting in >3-log fold (>99.9%) reduction in pathogen load in host cells (Fig. 5g-l). We also evaluated the spray characteristics of PCANS sprayed through a standard and commercially used VP3 multi-dose nasal spray pump (Aptar, USA). The droplet distribution data showed that 10% of PCANS droplets had size >10 µm, and 90% had size <200 µm (Fig. 5m), which is desirable to maximize the deposition in nasal cavity, while minimizing deposition into deep lungs. PCANS resulted in a wide plume angle within the ideal range of 35-55°, an ovality close to 1, covering a circular area of up to 8%, which is in line with the commercial nasal sprays (*53*, *54*)(Fig. 5m).

**Fig. 5:**
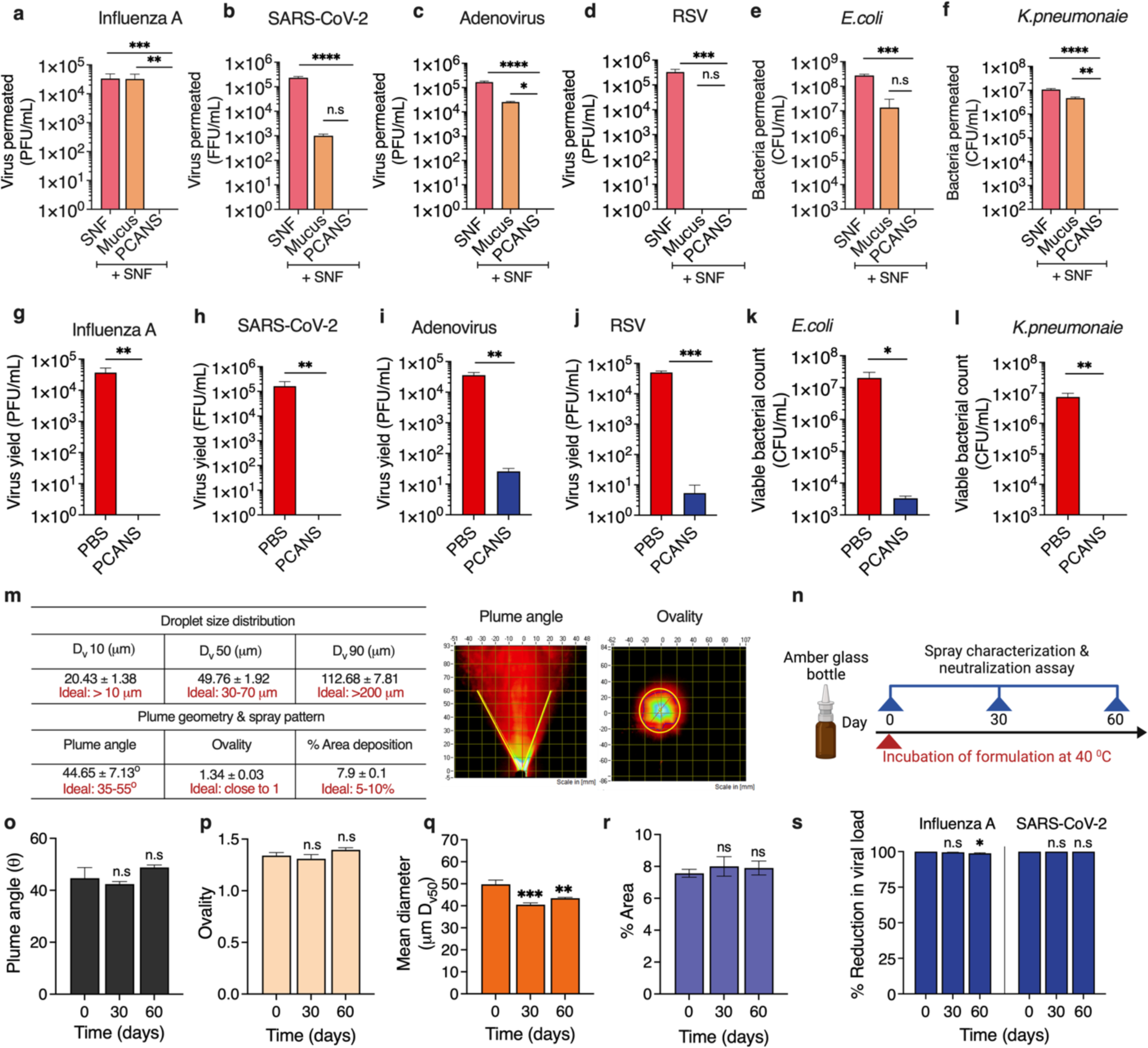
PCANS exhibits broad-spectrum physical barrier property with pathogen neutralization, and is sprayable and shelf-stable. **(a-b)** Amount of different viruses that permeated within 4 h through a simulated nasal fluid (SNF)-coated cell strainer or a cell strainer coated with simulated mucus/SNF mixture or PCANS/SNF mixture. Virus permeation was quantified by plaque assay in MDCK cells (IAV), Vero E6 cells (SARS-CoV-2), and Hep-2 cells (RSV and adenovirus). \*\*\*\**P* < 0.0001, \*\*\**P* < 0.001, \*\**P* < 0.01, \**P* < 0.05. n.s, not significant. Amount of **(e)** *E. coli* and **(f)** *K. pneumoniae* that permeated within 4 h through an SNF-coated cell strainer or a cell strainer coated with simulated mucus/SNF mixture or PCANS/SNF mixture. Bacterial permeation was quantified using a CFU plate count method. \*\*\*\**P* < 0.0001, \*\*\**P* < 0.001, \*\**P* < 0.01. n.s, not significant. **(g-k)** Viral titer for IAV, SARS-CoV-2, adenovirus, and RSV after treatment with PBS or PCANS. IAV and SARS-CoV-2 were incubated with PCANS for 10 min, while adenovirus and RSV were treated for 30 min. Anti-bacterial activity of PCANS against **(k)** *E. coli* and **(l)** *Klebsiella pneumoniae* using CFU plate count method after 30 and 10 min incubation, respectively. \*\*\**P* < 0.001, \*\**P* < 0.01, \**P* < 0.05. **(m)** Spray characteristics of PCANS. The droplet size distribution of PCANS was analyzed using a laser diffraction system. Representative images of single time delay plume angle and ovality ratio, captured using a high-speed digital camera and laser light sheet. **(n)** Experimental design to assess the stability of PCANS in accelerated temperature conditions (40°C). PCANS was stored in glass amber bottles. Aliquots were taken at different time points to investigate spray characteristics and pathogen neutralization efficacy. **(o)** Plume angle, **(p)** ovality, **(q)** mean droplet diameter and **(r)** spray deposition area over a period of 60 days. \*\*\**P* < 0.001, \*\**P* < 0.01 compared to day 0, n.s, not significant. **(s)** Percent reduction in the viral load of IAV and SARS-CoV-2 in their respective host cells after 10 min incubation with PCANS aliquoted at different time points in the stability study. \**P* < 0.05 compared to day 0. n.s, not significant. For **a-f** and **o-s**, *P* values were determined using one-way ANOVA with Tukey’s post-hoc analysis. For **g-l**, *P* values were determined using a two-tailed t-test. Data are presented as Means ± SD of biological repeats (n = 3, each experiment performed at least twice).

Shelf-stability is a key attribute governing the translational potential of formulations. We tested shelf-stability of PCANS over 60 days at 40°C temperature, as per the International Conference on Harmonisation (ICH) guidelines for stability testing under accelerated storage conditions (Fig. 5n). Over a period of 60 days, we observed no substantial variations in the spray characteristics, including plume angle, ovality, coverage area, and droplet size distribution (Fig. 5o-r and Fig. S10). PCANS also displayed no changes in its neuralization activity over 60 days, resulting in >99.99% reduction in Influenza A and SARS-CoV-2 viral loads in the host cells upon 10 min of incubation (Fig. 5s). Collectively, these data confirm the shelf-stability of PCANS.

### PCANS exhibits prophylactic activity *in vivo*

Next, in a proof-of-concept study, we investigated the prophylactic efficacy of PCANS against respiratory infection *in vivo*. PR8, a mouse-adapted strain of H1N1 Influenza virus, was used to induce infection. PR8 is a highly virulent strain that induces severe respiratory infection in mice(*55*), and can be lethal at a dose of 10 PFU(*56*). *In vitro* assay revealed excellent potency of PCANS to neutralize 10^6^ PFU of PR8 within 10 min of incubation, resulting in >5-log fold (>99.99%) reduction of the viral load in host cells (Fig. S11). To demonstrate efficacy *in vivo*, PCANS or PBS (10 µl) was administered prophylactically to both the nostrils of healthy mice on day 0 (Fig. 6a). Fifteen minutes later, animals were challenged intranasally with PR8 (250 PFU), a dose that been previously used by other groups(*57*, *58*). Remarkably, all mice in the PCANS-treated group survived for at least 10 days after the infection, whereas the PBS-treated group showed 100% lethality by day 8 (Fig. 6b). Over 10 days, no discernible change was observed in the body weight of the PCANS-treated animals, while significant weight loss was observed for PBS-treated ones after 3 days post-infection (Fig. 6c). PCANS also curtailed the lung viral titer to undetectable levels on days 2 and 4 post-infection, resulting in >5-log fold (>99.99 %) reduction compared to PBS-treated mice (Fig. 6d-e). Compared to healthy mice, mice infected with PR8 and treated with PBS showed significant differences in the levels of inflammatory cells, including leukocytes, neutrophils, lymphocytes, and macrophages in bronchoalveolar lavage (BAL) fluid (Fig. 6f-i). Prophylactic treatment of mice with PCANS restored the levels of inflammatory cells in BALF to normal. Additionally, cytokine profile from lung homogenate showed a significant reduction of IL-6 and TNF-α levels in PCANS-treated mice, as compared to the PBS-treated group (Fig. 6j-l). No reduction was, however, observed in the levels of IL-1β. Histological examination of lung sections revealed a substantial reduction in leukocyte infiltrates in PCANS-treated mice, as compared to the PBS-treated group, which showed an abundant presence of bronchial and alveolar infiltrates (Fig. 6m). Overall, compared to PBS-treated mice, we observed a significant reduction in pulmonary inflammation score for PCANS-treated group (Fig. S12). An escalated dose challenge was performed to determine the potency of PCANS to neutralize a higher viral load of PR8 (500 PFU). Compared to the PBS-treated group, prophylactic treatment with PCANS significantly improved survival and body weight and reduced lung viral titer on days 2 and 4 post-infection (Fig. S13). These data clearly indicate the potential of nasally administered PCANS to protect against respiratory infection in mice.

**Fig. 6:**
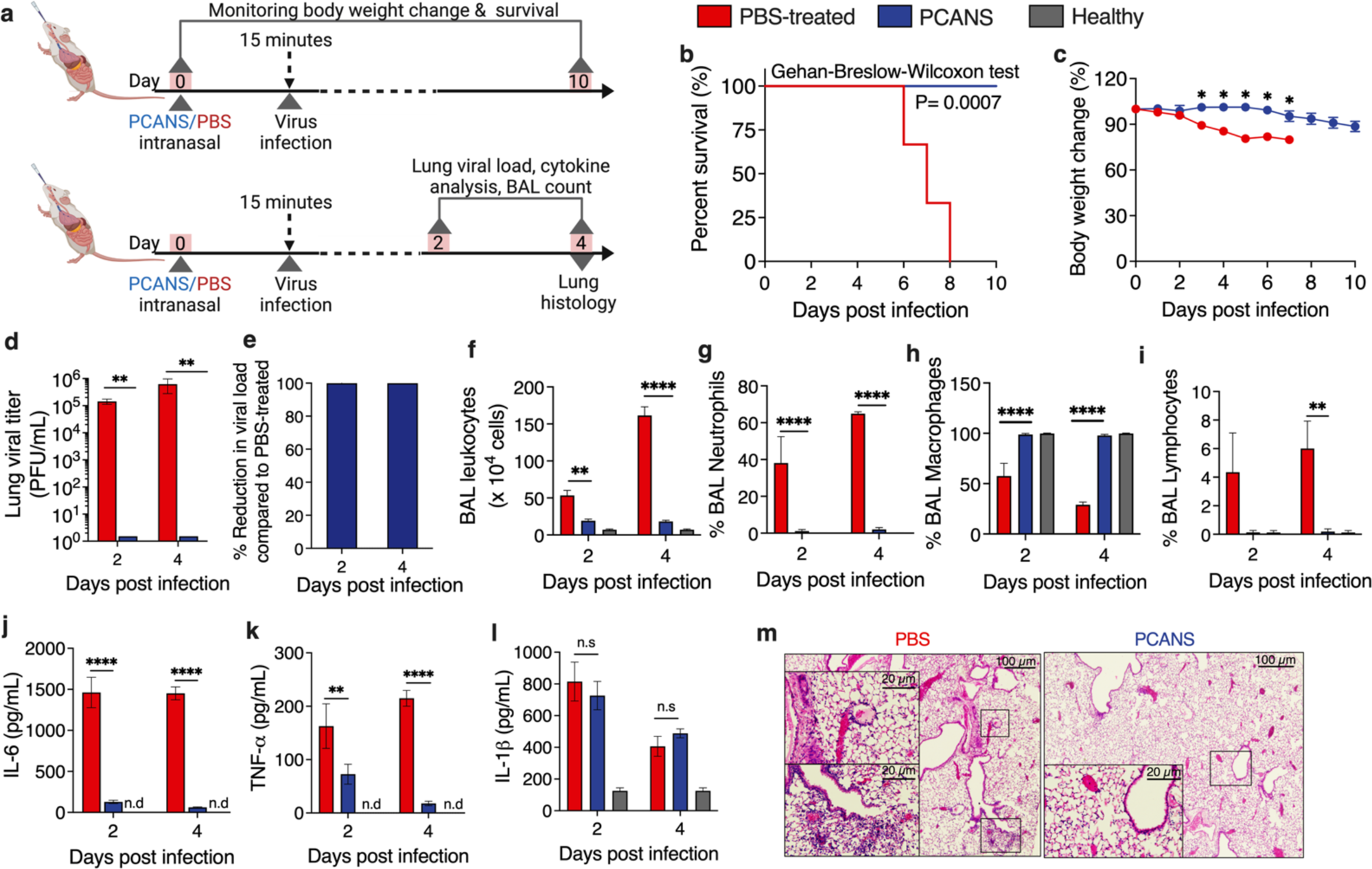
Pre-exposure prophylactic treatment with intranasal PCANS reduces respiratory infection in mice. **(a)** Experimental outline for the prophylactic efficacy study. C57Bl/6 mice received a single dose (10 µl) of PCANS or PBS before 15 minutes of intranasal inoculation with 250 PFU Influenza A/PR/8/34. One cohort of animals was followed for body weight changes and survival for a period of 10 days. Animals from a second cohort were euthanized on day 2 or 4 after infection to enumerate lung viral titer, inflammatory cell count in bronchoalveolar lavage (BAL) fluid, and inflammatory cytokine levels in lung homogenate. Hematoxylin and eosin (H&E) stained lung tissue sections from animals euthanized were assessed for inflammation. **(b)** Survival and **(c)** body weight change of mice over a period of 10 days post-infection. *P* = 0.0007 compared to the PBS-treated group for Kaplan-Meier survival curve. * *P* < 0.01 compared to PBS-treated group for body weight change curves. **(d)** Viral titer from lung homogenate of mice and **(e)** percentage reduction in viral load in the lungs on day 2 and 4 post-infection, as quantified by plaque assay performed in MDCK cells. \*\**P* = 0.001. **(f-i)** Inflammatory cell count in BAL on day 2 and 4 after infection. \*\*\*\**P* < 0.0001, \*\**P* < 0.01. Levels of **(j)** IL-6, **(k)** TNF-α and **(l)** IL-1β in lung tissues. \*\*\*\**P* < 0.0001, \*\**P* = 0.005. n.s, non-significant, n.d, not detected. **(m)** Representative images of H&E-stained lung tissue sections of virus-challenged mice that were prophylactically treated with PBS or PCANS. Histology images were captured using 10X and 40X objectives. Scale bar: 100 µm. High-magnified insets depict the difference in the extent of inflammatory infiltrates. Scale bar: 20 µm. For **b**, *P* values were determined using the Gehan-Breslow-Wilcoxon test. For **c**, *P* values were determined using one-way ANOVA with Brown-Forsythe. For **d** and **f-l**, *P* values were determined using two-way ANOVA with Tukey’s post hoc analysis. For o, *P* values were determined using one-way ANOVA with Tukey’s post hoc analysis. n=6 mice/group for **b**. Data in **c** are presented as Means ± SEM (n=6 mice/group). Data in **d-l** are presented as Means ± SEM (n=4 mice/group).

## Discussion

We report a chemoprophylactic nasal spray, PCANS – a radically simple and scalable pre-exposure prophylaxis approach to offer protection against current and emerging respiratory pathogens. PCANS acts *via* a multi-pronged approach that involves enhancing the capture of pathogen-laden large respiratory droplets from inhaled air, acting as a physical barrier to prevent the transport of respiratory pathogens through nasal lining, and rapid neutralization of a broad spectrum of respiratory pathogens. Unlike vaccines, which are pathogen-specific and exhibit reduced efficacy as the pathogen mutates(*59*), PCANS has broad spectrum activity, with potential to target both current and emerging respiratory pathogens. In a proof-of-concept study performed in mice, a single intranasal dose of PCANS was effective as early as 15 min after administration and provided protection against supra-lethal dosages of a highly virulent mouse-adapted strain of H1N1 Influenza virus (PR8), efficiently reducing the lung viral titer by over 80-99.99%. This implies potential use of PCANS as an additional layer of protection in conjunction with vaccines to minimize pathogen load, which is otherwise difficult to achieve with vaccines alone. For example, in a clinical study, participants vaccinated with BNT162b2 and mRNA-1273 had only 40 percent less detectable virus compared to those who were unvaccinated when infected(*60*).

PCANS embodies multiple advantages over previously developed chemoprophylactic nasal sprays. First, PCANS exhibits neutralization activity against multiple pathogens, including both bacteria and viruses (enveloped and non-enveloped). Most of the previously developed nasal sprays, on the contrary, consist of a single active ingredient that targets a limited type/class of pathogen. For example, nasal sprays containing iota and kappa carrageenan (Carragelose®/Dual Defence®), xylitol (Xlear Nasal Spray®), or ethyl lauroyl arginine hydrochloride (COVIXYL-V®) have reported activity only against IAV and SARS-CoV-2(*18*, *25–28*). Second, compared to other previously developed chemoprophylactic nasal sprays that require 0.5-2 h for pathogen neutralization(*25*, *28*), PCANS can rapidly neutralize both bacteria and viruses within 10-30 min, and also showed higher effectiveness. For instance, PCANS achieved a 5-log fold reduction in the viral load of SARS-CoV-2 within 10 min, while a xylitol-based nasal spray reduced viral titer by 2.5 log fold in 25 min(*61*). In this study, iota + kappa carrageenan, the key component of a commercially available nasal spray-Dual Defence, resulted in only 1-log fold reduction in the viral loads of both IAV and SARS-CoV-2 within 10 min, which is consistent with a previous report, where iota carrageenan reduced IAV load by 40% within 10 min(*21*). On the other hand, PCANS led to a 99.99% reduction (4-log fold) reduction in the viral load of both IAV and SARS-CoV-2 within the same contact time. PCANS also demonstrated excellent prophylactic efficacy in mice challenged with a supra-lethal dose of PR8 virus, resulting in >5-log fold reduction in the lung viral titer, and 100% survival of animals. Previous nasal sprays, on the other hand, have demonstrated sub-optimal efficacy. For example, a carrageenan-based nasal formulation only showed 60% survival of mice challenged with the PR8 virus(*21*). Third, the “drug-free” nature of PCANS is favorable for the regulatory process, which could be tedious for chemoprophylactic approaches based on investigational new drugs such as IgM-14(*62*). Also, since all the components used in PCANS are commercially available off-the-shelf and require simple mixing without chemical modifications, our approach is amenable to scale-up and large-scale manufacturing. Fourth, Chemoprophylactic nasal sprays developed in the past have typically depended on a single mode of action, which involves either neutralizing pathogens or preventing their contact or entry into the nasal lining. We believe that this singular approach has, in part, contributed to their observed sub-optimal efficacy in clinical settings. For example, in a clinical study, a xylitol-based nasal spray resulted in only 62% fewer infections when compared to placebo infected with SARS-CoV-2(*23*). Similarly, a 5B5 monoclonal antibody-based nasal spray only showed protective effectiveness of 60% and 20%, against delta and omicron variants of SARS-CoV-2. PCANS is equipped with three critical attributes that involve capturing pathogen-laden large respiratory droplets from inhaled air, acting as a physical barrier to prevent the transport of pathogens into the nasal lining, and rapid neutralization ability against a broad spectrum of pathogens. Fifth, PCANS is safe for daily administration, as demonstrated in mice, which is a significant advantage over previously developed povidone iodine-based anti-viral nasal sprays(*19*, *30*), which are associated with iodine burns, thyroid toxicity, and disruption of the mucosal barrier, constraining repeated administration. Similarly, frequent use of a nitric oxide (NO)-inducing nasal spray (SaNOtize®), which has shown potential in post-exposure prophylaxis of SARS-CoV-2 infection, can result in elevated Th2 cytokines, which mediate autoimmune disorders(*63*). In addition, excessive NO can cause tissue damage and cell death(*64*). The incidence of such adverse effects with PCANS is likely to be low, as the formulation is devoid of immunomodulatory molecules such as NO and steroids. Finally, PCANS exhibits an excellent residence time of 8 h in the mouse nasal cavity and provides protection for several hours. To the best of our knowledge, this is the longest nasal residence time that has been reported for chemoprophylactic nasal sprays in mice. Such a long nasal residence time would potentially minimize dosage frequency in humans, offering an advantage over previously developed chemoprophylactic approaches, including SaNOtize® that require 3-6 doses per day due to short half-life of NO(*64*, *65*). Mechanistic studies performed in mice demonstrated that the prolonged nasal residence time of PCANS is due to the presence of tween-80. Although different surfactants, including tween-80 have previously been shown to slow mucociliary clearance by reducing the cilia beat frequency(*52*), further study is warranted in the future to comprehend the mechanism of tween-80-induced delayed nasal clearance of PCANS. However, our study confirmed that daily administration of PCANS in mice is safe, with no evidence of inflammation or any other nasal toxicity observed after a repeated dosing for at least 14 days.

Our study has several strengths. First, the components constituting PCANS were identified *via* rigorous *in vitro* and *in vivo* screenings of multiple excipients from the IID and GRAS list of the FDA, and their different concentrations and combinations. These extensive screening experiments were aimed to optimize the key parameters, including sprayability, mucoadhesiveness, capture of respiratory droplets, physical barrier property, broad spectrum pathogen neutralization activity, and nasal residence time. Second, to evaluate the respiratory droplet capturing ability of PCANS, we used a 3D-model of human nasal cavity (Koken cast), which is based on a female cadaver and has been previously used for *in vitro* evaluation of nasal drug delivery, as it replicates all the anatomical intricacies of human nasal cavity(*51*). Third, physical barrier property of different biopolymers was evaluated by two complementary techniques – quantification of viral transport using a plaque-forming assay and quantification of the transport of small molecule dye using fluorescence spectroscopy. Fourth, we demonstrated broad spectrum physical barrier property and neutralization ability of PCANS in five different pathogens – three enveloped viruses (IAV, SARS-CoV-2, RSV), one non-enveloped virus (adenovirus), and two bacteria (*E. coli* and *K. pneumoniae*). To our understanding, this is the first report demonstrating such a broad-spectrum activity of a chemoprophylactic nasal spray. Lastly, to demonstrate prophylactic efficacy of PCANS *in vivo*, we used a highly virulent mouse-adapted strain of H1N1 Influenza virus (PR8) that induces severe respiratory infections in mice(*55*). Prophylactic efficacy of PCANS was demonstrated against three different dosages of the virus, which were 10-50 times higher than the previously established lethal dose for PR8 in mice(*56*). Due to its prolonged nasal residence time, PCANS was effective for several hours after nasal administration.

In conclusion, PCANS presents a promising chemoprophylactic approach against respiratory infections. Besides its potential to act as a first line of defense against respiratory pathogens and emerging variants for which there are no vaccines available, our approach could also be used as an added layer of protection with existing vaccines. Given its broad-spectrum prophylactic activity and excellent shelf stability, we anticipate PCANS holds the potential for global distribution, especially in countries with low vaccination rates against respiratory pathogens. Alongside, the benefits of PCANS can also be extended to immunocompromised patients, high-risk individuals with co-morbidities, and vaccine-hesitant populations. Its pocket-sized spray format allows for easy portability, making it convenient to carry during social gatherings and travel. With these significant benefits, we believe PCANS will experience rapid widespread adoption, enhancing the accessibility of respiratory infection prevention. By enabling people to breathe clean and minimizing the transmission of respiratory infections, PCANS can play a pivotal role in safeguarding public health worldwide.

## Methods

### Preparation of biopolymer solutions and PCANS

Biopolymer solutions were prepared by the addition of the biopolymer (0.2 to 2% w/v) to ultrapure deionized sterile water (Invitrogen). The solution was then mixed to attain a homogenous mixture with slight heating at 60°C. Biopolymers including gellan (Gelzan), pectin, carboxymethylcellulose (CMC), hydroxypropyl methylcellulose (HPMC), carrageenan, xanthan gum, and Carbopol were purchased from Sigma Aldrich. To prepare PCANS, gellan and pectin solutions were mixed in a ratio of 1:1, followed by the addition of tween-80 (Sigma Aldrich). The solution was then supplemented with benzalkonium chloride (BKC) (Sigma Aldrich) and subjected to immediate mixing by pipetting up and down several times. Finally, phenethyl alcohol (Sigma Aldrich) was added, and the pH of the solution was adjusted to 5.5. For cell culture experiments and *in vivo* efficacy study, the individual components of PCANS were sterile filtered using 0.2 μm PVDF syringe filters (EMD Millipore) and combined as described above.

### Preparation of simulated nasal fluid (SNF) and simulated mucus

SNF was prepared by dissolving 1.32 g sodium chloride (150 mM), 447 mg potassium chloride (39.9 mM), and 88.5 mg calcium chloride (5.3 mM) in 150 mL ultrapure deionized sterile water and filtered using 0.2μm filter(*68*). The healthy simulated mucus was formulated by dissolving 0.6 mg mucin from porcine stomach Type II (Sigma Aldrich), 0.8 mg mucin from porcine stomach Type III (Sigma Aldrich), 0.32 mg bovine serum albumin (Sigma Aldrich) in 10 mL ultrapure deionized water containing 20 mM HEPES buffer and 38 mM sodium chloride solution(*69*). The mixture was stirred vigorously under slight heating to attain a homogenous solution.

### Rheological measurements

Dynamic viscosity behavior of biopolymer solutions was evaluated using a rotational rheometer (Discovery HR-2, TA Instruments) using a 40 mm diameter cone with a geometry angle of 1^0^. Samples were subjected to a linear shear rate ramp up to 40 s^−1^ at 25° C to mimic the strain encountered by the formulation when actuated through the nozzle of the spray device. The viscosity of the biopolymer solution was measured during the upward ramp in triplicates. The sol-gel transition of biopolymer solutions with and without the presence of SNF was evaluated by rotational rheology. The mechanical strength in terms of storage modulus was assessed by applying amplitude sweep with a varying oscillatory strain at 1 Hz at 37° C.

### *Ex vivo* mucosal retention study

Tissue harvested from sheep was cut open to expose the mucosal surface and trimmed down to 75×26 mm. Mucosal tissue was then mounted on a glass slide facing upwards and positioned at 45^0^ to align it with the spray actuation angle. The tissue was initially moistened with SNF using a generic nasal spray device, and excessive fluid was removed with sterile wipes. Brilliant green dye (Sigma Aldrich) loaded polymeric solution was sprayed, keeping the spray nozzle tip at a distance of 5 cm from the slide surface. The slides were examined for runoff/drip after 4 h of spraying. The distance traveled by the polymer solution down the glass slide from the bottom end of formulation deposited on mucosal tissue was measured as drip length. Drip length of free dye was considered 100%.

### Cell culture

Madin-Darby canine kidney cells (ATCC®) were cultured in T-175 flasks (CELLTREAT) at 37°C and 5% CO_2_ in DMEM (Gibco) supplemented with 10% fetal bovine serum (FBS) (Gibco) and 1% penicillin-(streptomycin (Invitrogen). Hep2 cells and Vero E6 cells (ATCC®) were cultured at Integrated Biotherapeutics (IBT) Bioservices in T-75 flasks at 37°C and 5% CO_2_ in EMEM supplemented with 10% FBS and 1% penicillin-streptomycin. Human nasal epithelial cells (ATCC) were cultured in T-175 flasks at 37°C and 5% CO_2_ in EMEM supplemented with 10% FBS and 1% penicillin-streptomycin.

### Production of NanoLuc Luciferase expressing recombinant SARS-CoV-2

All replication-competent SARS-CoV-2 experiments were performed in a BSL-3 facility at the Boston University National Emerging Infectious Diseases Laboratories. A recombinant SARS-CoV-2 virus expressing a NeonGreen fluorescent protein (rSARS-CoV-2 mNG)(*70*) was generously provided by the Laboratory of Pei-Yong Shei. To propagate the virus, 1×10^7^ Vero E6 cells were seeded in a T-175 flask one day prior to propagation. The next day, 10 µL of rSARS-CoV-2 mNG virus stock was diluted in 10 mL of OptiMEM, added to cells, and then incubated for 1 h at 37°C. After incubation, 15 mL of DMEM containing 10% FBS and 1% penicillin/streptomycin was added to cells. The next morning, media was removed, cells were washed with 1X PBS and 25 mL of fresh DMEM containing 2% FBS was added. Virus was incubated for an additional 48 h. The supernatant was collected at 72 h, filtered through a 0.22 µm filter, and stored at −80°C. The viral stock was thawed and concentrated by ultracentrifugation (Beckman Coulter Optima L-100k; SW32 Ti rotor) on a 20% sucrose cushion (Sigma-Aldrich, St. Louis, MO) at 25,000 x g for 2 h at 4°C. Media and sucrose were then discarded, pellets were dried for 5 min at room temperature, and viral pellets were resuspended in 100 µL of cold 1X PBS at 4°C overnight. The next day, concentrated virus was combined, aliquoted and stored at −-80°C.

### *In vitro* physical barrier assay

A 70 μm pore size mesh cell strainer was coated with 15 µL of mucus, or a biopolymer solution, or PCANS. The formulation was spread evenly using a sterile stainless-steel spatula with a tapered end. To facilitate *in situ* gelation, 15 µL of SNF was added, covering the entire surface of the strainer. The strainer was placed in a 6-well plate containing 0.9 mL of serum-free DMEM (for virus/bacteria penetration) or ultrapure deionized water (for rhodamine B isothiocyanate penetration) in each well, and 0.1 mL of diluted virus (∼1 x 10^5^ PFU/mL)/bacteria (1×10^7^ CFU/mL) stock or rhodamine B isothiocyanate (1mg/mL) was added to the upper compartment of the strainer. After 4 h of incubation at 37°C, medium or deionized water from the bottom reservoir was retrieved, and the viral titer permeated through the hydrogel layer was quantified using plaque assay for IAV performed in MDCK cells, crystal violet staining for RSV performed in Hep-2 cells, immunostaining for adenovirus performed in Vero E6 cells, focus forming assay for SARS-CoV-2 in Vero E6 cells, and colony forming unit (CFU) plate count method for bacteria, as described in the following sections. The permeation of dye through biopolymer solution/mucus was quantified by measuring the fluorescence intensity using a microplate reader.

### *In vitro* neutralization assay with Influenza A

Neutralization activity of different excipients and PCANS was evaluated by plaque assay. MDCK cells were seeded at a density of 2-3 million cells per well in a 6-well plate and then incubated at 37°C to achieve ∼80-90% confluency one day before infection. On the day of infection, 50 µL of HKx31 Influenza A virus (H3N2, 5×10^4^-1×10^5^ PFU/mL) (BEI Resources) in infection media (serum-free DMEM containing 3 μg/mL TPCK-trypsin) was pre-treated with 50 µL of PCANS, biopolymer solution, surfactant solution, alcohol solution or PBS. Samples were vortexed for 10 seconds and incubated at 37°C for 10 or 60 min. After incubation, samples were centrifuged for 1 min at 1000 RPM, and the supernatant was subjected to a 10-fold serial dilution until eighth dilution using infection medium. MDCK cells were then exposed to pre-treated virus dilutions for 1 h. After infection, an overlay growth medium containing 2X DMEM with 2% agarose (50:50) was poured onto the top of the cell monolayer and incubated for 72 h. The overlay was removed, and cells were then fixed using 1 mL of 10% formalin and left for 1 h at room temperature, followed by the addition of 1% crystal violet for 5-15 min. Wells were washed with water and left to dry out and PFUs were counted to determine the viral titer.

### *In vitro* neutralization assay with SARS-CoV-2

The day prior to infection experiment, 8×10^4^ Vero E6 cells/well were plated in a 24-well plate. To perform neutralization assay, 50 µL of PCANS, biopolymer solution, surfactant solution, alcohol solution or PBS was mixed with 8×10^4^ PFU of SARS-CoV-2 mNG in 50 uL of infection media (OptiMEM (Gibco) containing 3 μg/mL TPCK-trypsin), vortexed and centrifuged briefly prior to incubation at 37°C for 10 or 60 min. After incubation, samples were centrifuged for 1 min at 1000 RPM and serially diluted 10-fold until eighth dilution with infection medium. Of each dilution, 200 µL was then plated into a 24-well plate and incubated for 1 h at 37°C prior addition of 800 µL of 1.2% Avicel (Dupont). Following a 24 h incubation period at 37°C, Avicel was removed, cells were washed with 1X PBS and fixed for 3 h with 10% neutral buffered formalin. Focal forming units (FFU) per mL were determined by counting NeonGreen expressing foci using an Evos M5000 fluorescent microscope (Thermo Scientific).

### *In vitro* neutralization assay with adenovirus and respiratory syncytial virus

The broad-spectrum neutralization potency of PCANS was evaluated against adenovirus type 5 (ADV-5, ATCC, VR-2554™) and respiratory syncytial virus strain A2 (RSV-A2, ATCC, VR-1540™) using plaque assay. Briefly, the day prior to the infection, 1×10^5^ Vero E6 cells/well or 1.5×10^5^ Hep-2 cells/ well were plated in a 24-well plate for ADV-5 and RSV-A2, respectively. On the day of infection, 50 μL of PCANS was mixed with 50 μL of virus (1 x10^6^ PFU/mL of ADV-5 and 2×10^6^ PFU/mL of RSV-A2) in the infection media and incubated at 37°C for 30 min. The pre-treated mixture was 10-fold serially diluted in infection media after the incubation. Cells were washed with serum-free media before infection and 200 μL of each dilution was transferred to the cells for a 1 h incubation prior to the addition of a 1 mL overlay medium containing methylcellulose. Following a 72 h incubation, the overlay layer was removed, and cells were fixed using 10% formalin with subsequent immunostaining for Vero E6 cells and crystal violet staining for Hep-2 cells. Plaques were counted using a plaque reader (Bioreader-600-Vα).

### *In vitro* neutralization assay with bacteria

The neutralization potency of components and PCANS was studied against gram-negative bacteria including *Escherichia coli* (*E. coli*) and *K. pneumoniae.* An overnight culture of bacteria was prepared in 5 mL tryptic soy broth (TSB, Sigma Aldrich) media. On the day of the experiment, bacteria suspension was adjusted to obtain an OD_600nm_ = 0.2, which corresponds to 10^8^ CFU/mL. A 50 µL of bacterial suspension in TSB media was incubated with 50 µL of PCANS, biopolymer solution, surfactant solution or alcohol solution at 37°C for 10 or 60 min. After incubation, the sample/bacteria mixture was 10-fold serially diluted in 1X PBS, and 10 μL of each dilution was plated onto pre-poured LB (Luria Broth, HiMedia Laboratories Pvt Ltd) agar plates followed by an incubation of 16-18 h at 37°C, 5% CO_2_. The plates were then counted for CFUs.

### TEER assay and *in vitro* cytotoxicity of tween-80

RPMI 2650 cells were seeded on the apical part of Transwell inserts (6.5 mm polyester membrane ∼ 0.4 µm pore size, Corning) at a density of 1.5 x 10^5^ cells/cm^2^ in 0.1 mL EMEM. The basolateral compartment of the insert was filled with 0.6 mL EMEM media supplemented with 10% FBS followed by incubation at 37°C. On day 4, the medium was removed from the top of the inserts, and media volume in the bottom well was reduced to 200 µL. Every 2 days the medium was changed, and TEER was measured. An epithelial volt ohmmeter (World Precision Instrument) was used to measure the impedance. Until the monolayer formed with a constant impedance around 12, cells were grown with an air-liquid interface. On day 12, TEER was measured prior to the treatment of cells with surfactants. 200 µL of medium containing Triton X-100 (0.1% w/v) or tween-80 at different concentrations was added to the insert. Plate was incubated at 37°C for 4 h. After incubation, wells were replenished with fresh medium, and TEER was measured after 4, 5, 12, and 24 h. The cytotoxic effect of tween-80 at different concentrations was also studied on RPMI 2650 cells. Briefly, 20,000 cells/well were seeded in a 96-well plate and incubated at 37°C overnight to achieve 70-80% confluency. Tween-80 solution in 0.2 mL EMEM media was added to the wells, followed by an incubation for 24 and 48 h. The metabolic activity of RPMI 2650 cells was measured using an XTT (2,3-bis(2-methoxy-4-nitro-5-sulfophenyl)-2H-tetrazolium-5-carboxanilide) assay kit (ATCC®) according to the manufacturer’s protocol.

### Capture of respiratory droplets

The inner surface of a glass twin impinger’s (Copley Scientific) oropharyngeal region (denoted by red arrows in Fig.4a) was coated with SNF followed by spraying the gellan and pectin mixture without or with different concentrations of tween-20, tween-80 or BKC using a VP3 nasal spray pump (Aptar). Droplets with mass median aerodynamic diameter >5 µm and laden with rhodamine B-loaded liposomes (size ∼400 nm) were generated using a jet nebulizer. Nebulized droplets were administered into the impinger under vacuum at a flow rate of 15 L/min for 3 min. The gel was retrieved, and fluorescence intensity was quantified at an excitation and emission wavelength of 543 and 580 nm. Rhodamine B-loaded liposomes were synthesized using the thin-film hydration method(*71*). Briefly, the lipids, DSPE-PEG (2000) amine (Avanti Polar lipids), cholesterol (Sigma) and L-α-phosphatidylcholine, hydrogenated (Soy) (HPC, Avanti Polar lipids) were dissolved in chloroform to prepare a 10mg/mL lipid stock solution in 1:1:3 molar ratio. A 2 mL of lipid stock solution was added to a round-bottom flask containing 0.8 mL of rhodamine B isothiocyanate from a 1mg/mL stock. The organic solvent was then evaporated using a rotary evaporator for 5 min to form a thin lipid layer. The lipid film was then hydrated using 10 mL ultrapure water (Invitrogen) and silica glass beads were added to the flask to suspend the lipid in the solution with vigorous shaking using the rotary evaporator at 40°C for 45 min. The hydrated lipid suspension was sonicated (Probe sonicator) at 30% amplitude for 1 min with a 2sec pulse on and off condition. The size of liposomes was then analyzed using a Zeta Analyzer (Malvern). To emulate the capture of pathogen-laden droplets in the human nasal cavity, a 3D transparent, silicone human nose model (Koken Co, Ltd) was used. The anterior region of the Koken model was deposited with SNF followed by the gellan and pectin mixture or PCANS with a single actuation using a nasal spray pump (Aptar). Koken model was connected to a vacuum pump at an air flow rate of 15 L/min and rhodamine B-loaded liposomes were then nebulized for 1 min. The model was disassembled to retrieve the formulation and captured dye-loaded droplets after nebulization. The capture of droplets was measured by quantifying the fluorescence intensity at an excitation and emission wavelength of 543 and 580 nm.

### Spray characterization

Multi-dose nasal spray vials were filled with water or gellan solution or PCANS. The pump (140 µL) with an insertion depth of 1.8 cm (Aptar) was used to study the spray characteristics including plume geometry, spray plume, and droplet size distribution. Three replicate measurements were performed for each sample. Plume geometry and spray pattern were measured using a Spray-View® measurement system (Proveris Scientific, Hudson, MA) at a distance of 30 mm from the nozzle orifice of the actuator. This acquisition system employs a high-speed digital camera and laser light sheet to capture images. Data were analyzed using an image processing software, Viota®. Actuation parameters including velocity, acceleration and hold time, and settings for camera and laser were kept identical across all the samples. Plume geometry measures the angle of plume ejected from the nozzle orifice. Ovality and plume area were evaluated to quantify the spray pattern of the samples. Ovality is defined as the ratio of maximum to minimum cross-sectional diameter of the spray plume. A uniform circular plume with an ovality close to 1 can be considered an optimal condition for nasal sprays.(*72*) Droplet size analysis of samples was inspected using a Malvern Spraytec® laser diffraction system. The FDA recommends reporting the measurements of size distribution data at D(v,0.1), D(v,0.5), and D(v,0.9) thresholds which correspond to the size of 10%, 50%, and 90% droplets by volume distribution, respectively(*53*). It is suggested to have droplet population with D(v,0.1) > 10 μm, D(v,0.5) between 30-70 μm and D(v,0.9) <200 μm. Droplet populations smaller than 10 μm have a propensity to induce a non-targeted deposition at the lungs, and droplets greater than 200 μm tend to drip/ run off the nasal cavity(*53*).

### Shelf-stability study

PCANS (15 mL) was filled in a sterile multi-dose nasal sprays (Aptar) capped with the actuator. The nasal spray vials were stored at an accelerated temperature condition (40° C). Aliquots were retrieved at different time points and evaluated for neutralization activity against IAV and SARS-CoV-2 using plaque forming and focus forming assays, respectively, as described above. Aliquots were collected from three different vials. Similarly, 5 mL aliquots were used to evaluate the spray features, including spray pattern, plume geometry, and droplet size distribution.

### Mice

Animal experiments were conducted according to ethical guidelines approved by the Institutional Animal Care and Use Committee (IACUC) of Brigham and Women’s Hospital. Experiments were conducted in 6-8 weeks-old C57BL/6 mice (Jackson Laboratories, USA). Mice were maintained under pathogen-free conditions and randomly assigned to various experiment groups, irrespective of gender. The group size of animals in experiments was decided based on the minimum number of animals required to attain a statistical significance of *P*<0.05 among different test groups. For mouse model of influenza infection, experiments were conducted in Biosafety Level 2 according to ethical guidelines approved by the Institutional Animal Care and Use Committee (IACUC) of Brigham and Women’s Hospital.

### *In vivo* biodistribution and nasal retention

Nasal retention of the formulation was performed in mice. Briefly, C57BL/6 mice were administered with 10 µL per nostril of free DiR (Thermofisher) or PCANS mixed with DiR at a final concentration of 10 µg/ml). Mice were euthanized at different time points and nasal cavity was harvested and imaged using IVIS (Bruker’s In-Vivo Xtreme optical and x-ray *in vivo* imaging system) at an excitation and emission wavelength of 680/700 nm. Vital organs such as lung, liver, spleen, kidney, and heart, were also imaged at 2 and 24 h time points. To determine the mechanism of long residence time, animals were intranasally instilled with DiR-mixed gellan and pectin mixture without or with BKC and tween-80. After 8 h, animals were euthanized to harvest and image the nasal cavity using Perkin Elmer IVIS Lumina II and the total flux was expressed in (p/sec/m^2^/sr).

### *In vivo* prophylactic activity of PCANS

Mice were intranasally instilled with 10 μL PCANS or PBS into each nostril under brief anesthesia using isoflurane. After 15 min, animals were challenged with 250 or 500 PFU of PR8 intranasally. One cohort of animals was followed for body weight changes and survival for a period of 10 days. Animals were euthanized when the body weight was reduced to 20%. Animals from a second cohort were euthanized either on day 2 or 4 after infection to enumerate lung viral titer, inflammatory cell count in bronchoalveolar lavage (BAL) fluid, and inflammatory cytokine levels in lung homogenate. BAL fluid was isolated by gently instilling saline solution into bronchioles with a catheter inserted through the trachea. The total cells and immune cell types from the collected BAL fluid were quantified using Diff-quik kit as per manufacturer’s protocol. For lung viral titer and cytokine profiling, left lung was homogenized and centrifuged at 2000 g for 10 min at 4°C to collect the supernatant. The obtained supernatant was further used for downstream assays. Viral titer was enumerated using plaque assay with MDCK cells, as detailed above. Cytokine profiling was performed using respective ELISA kits of IL-6, TNF-α, and IL-1β (BioLegend®) according to the manufacturer’s protocol. Histopathology of the right lung was determined using hematoxylin and eosin staining, and inflammation scoring was performed as reported previously(*73*).

### Statistics

Statistical analysis and graphing were conducted using Graphpad Prism. A one-way ANOVA with Tukey’s post hoc analysis was used to compare multiple groups. Two-way ANOVA with Tukey’s multiple comparison tests was used to analyze the data with two variables. To evaluate the efficiency of PCANS, survival plots were generated using the Kaplan-Meier survival curve, and the statistical significance of the results was analyzed using the Gehan-Breslow-Wilcoxon test. *P* values for the body weight changes were determined using one-way ANOVA with Brown-Forsythe post hoc analysis. A *P*-value of less than 0.05 was considered statistically significant.

## Acknowledgments

We acknowledge funding support from Gillian Reny Stepping Strong Center for Trauma Innovation at the Brigham and Women’s Hospital (to NJ and JMK), Startup Funds from the Department of Anesthesiology, Perioperative, and Pain Medicine at the Brigham and Women’s Hospital (to NJ), Fulbright-Nehru Postdoctoral Fellowship (to JJ), and Boston University (to FD) and the Peter Paul Career Development Award (to FD). The metered dose spray pumps were generously gifted by Aptar Inc. We acknowledge Integrated BioTherapeutics (IBT) Bioservices for evaluating the neutralization activity of PCANS against RSV and adenovirus, and Proveris Scientific for characterizing the spray patterns.

## Author contribution

Conceptualization: J.J., H.M.B., J.R.Q., Y.T., J.M.K., N.J. Data curation: J.J, H.M.B., J.R.Q., D.K., E.B. Data analysis: J.J., H.M.B., J.R.Q., D.K., E.B. Funding acquisition: J.M.K., N.J. Investigation: J.J., H.M.B., J.R.Q., Y.M., E.B., P.S., K.S., O.S., R.N., E.A., S.R., J.K. Methodology: J.J., H.M.B., J.R.Q., D.K., S.K., X.L.L., J.M., J.G., J.N.L, A.Y., F.D. Project Administration: Y.T., J.M.K., N.J. Supervision: F.D., Y.T., J.M.K., N.J. Validation: J.J., H.M.B., J.R.Q., J.M.K., N.J. Manuscript Writing - original draft: J.J., H.M.B., S.R., N.J., Manuscript Editing: J.J., H.M.B., D.K., F.D., Y.T., J.M.K, N.J.

