## Supplementary figures for "A Drug-Free Pathogen Capture and Neutralizing Nasal Spray to Prevent Emerging Respiratory Infections"

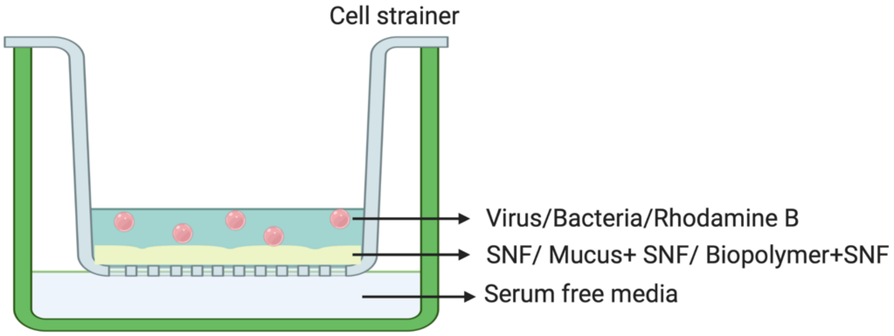


**Figure S1.** Experimental setup to evaluate physical barrier property of different mucoadhesive biopolymers. Biopolymers were coated on a cell strainer (pore size 70 μm) and permeation of Influenza A virus (IAV), bacterial or free rhodamine B dye was evaluated. The amount of virus, bacteria, or dye permeated was quantified from the acceptor compartment after 4 h.

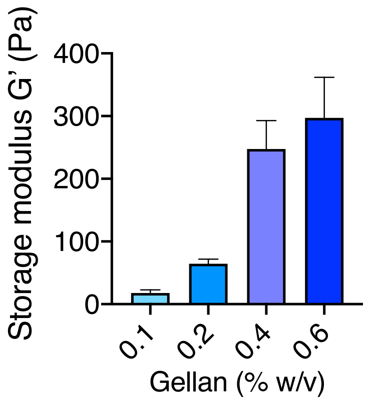


**Figure S2.** Concentration-dependent increase in storage modulus of gellan in the presence of SNF.

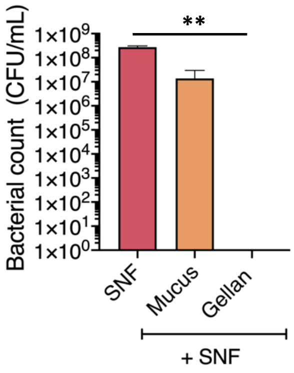


**Figure S3**. Inhibition of *E.coli* permeation by a physical barrier layer of nasal formulation imparted by crosslinked gellan (0.2% w/v) with SNF. **$P$ =0.0024. Data are presented as Means ± SD. $P$ values were determined using Tukey's post hoc analysis by one-way ANOVA.

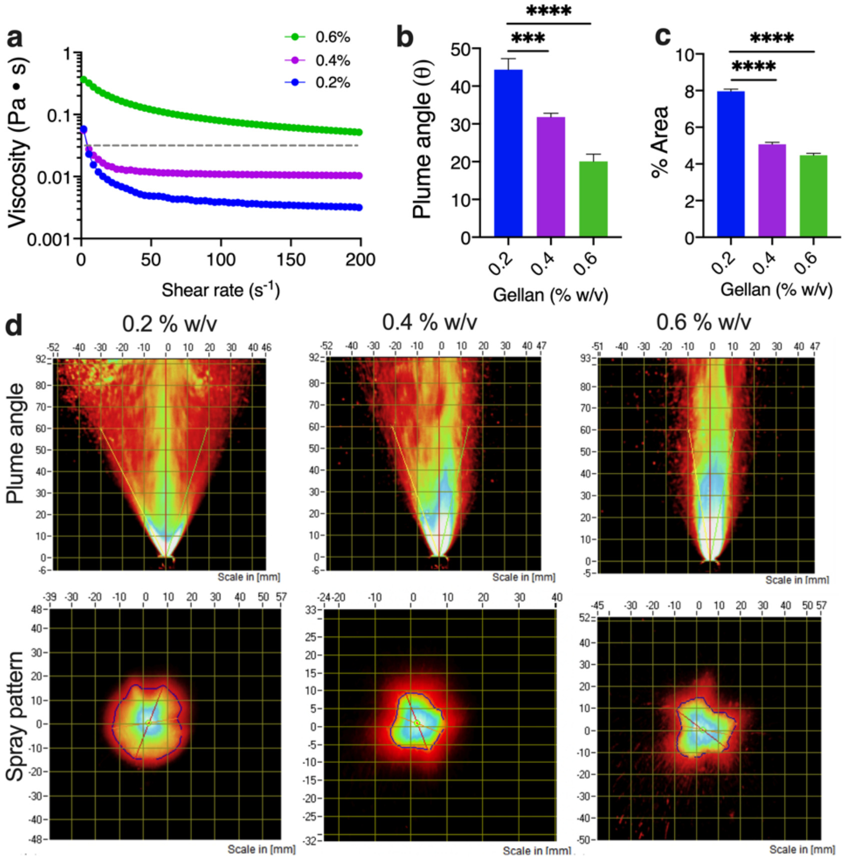


**Figure S4.** Concentration-dependent effect of gellan on viscosity and spray characteristics. a) Viscosity measured as a function of shear rate up to 200 $s^{-1}$ at 25°C for different concentrations of gellan. Quantitative measure of b) plume angle and c) spray coverage area was conducted to identify the concentration of gellan with maximum coverage area. ****$P$ < 0.0001,***$P$ = 0.0008,**$P$ = 0.0011 d) Representative images of plume angle and spray pattern. $P$ values were determined by one-way ANOVA using Tukey's post hoc analysis

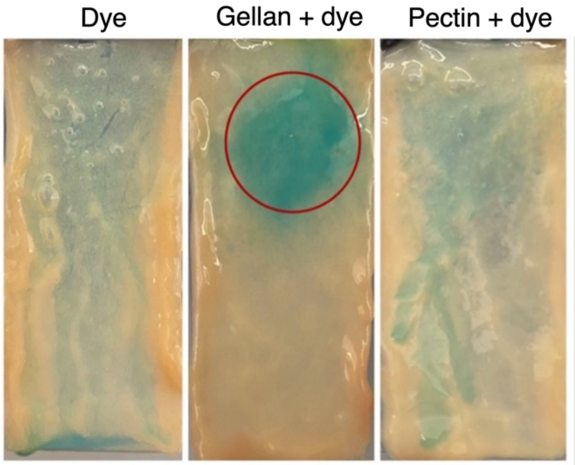


**Figure S5.** Drip length of mucoadhesive polymers on intestinal mucosa from pigs. Free dye or dye mixed with gellan and pectin was sprayed on mucosal tissue. The red circle denotes the initial spray area.

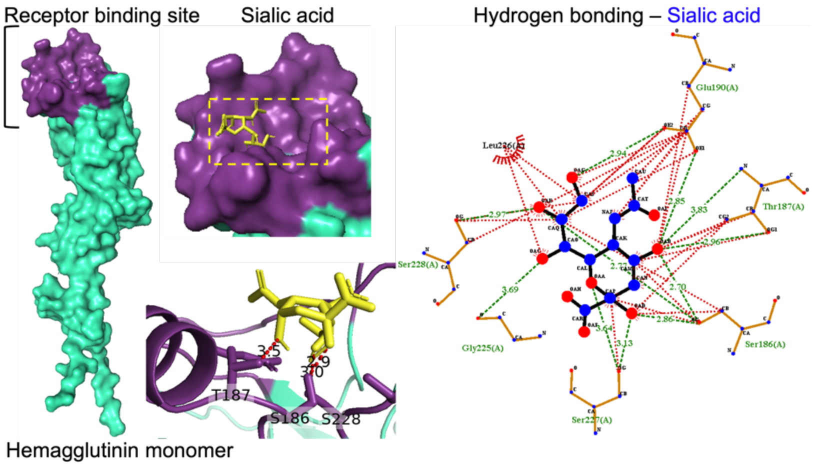


**Figure S6.** Interaction analysis of sialic acid with receptor binding domain of influenza A virus. Sialic acid (colored in yellow) binds to receptor binding site of hemagglutinin monomer (colored in violet) through hydrophobic interactions with Ser227 and Glu190, and hydrogen bonding with Ser288, Ser186, and Thr187.

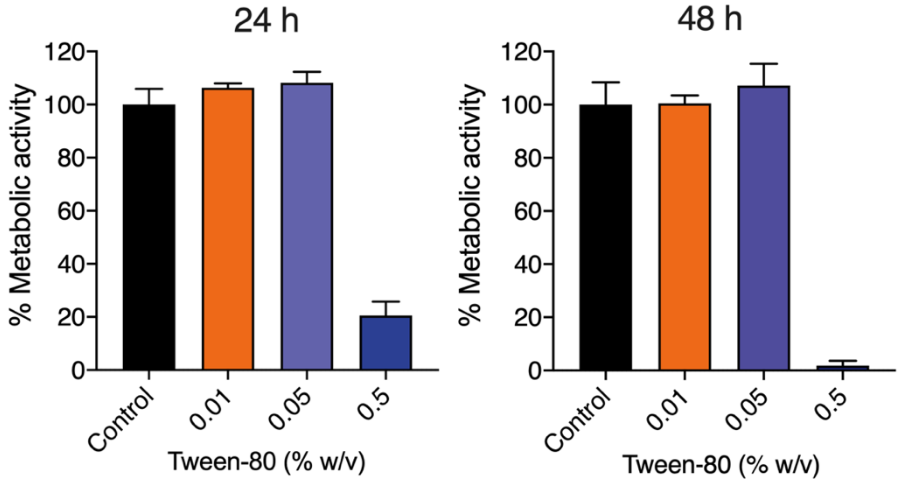


**Figure S7**. Influence of tween-80 on percentage cell viability of human nasal epithelium (RPMI-2650) cells after 24 and 48 h of incubation.

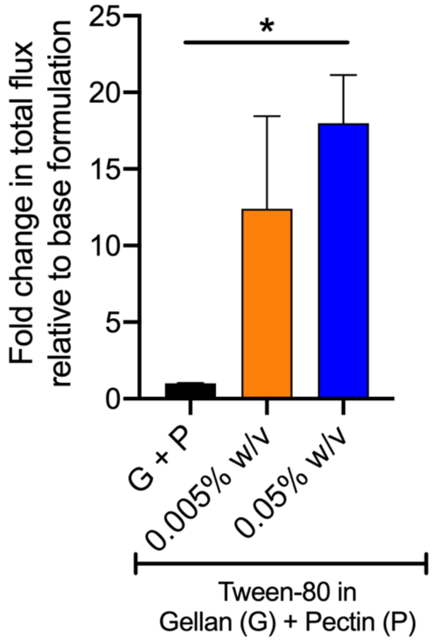


**Figure S8. Nasal retention of formulation loaded with NIR dye**. Concentration-dependent influence of tween-80 on the residence time of nasal formulation constituted of gellan and pectin after 8 hours of administration. ,*P=0.0392, *P* values were determined by one-way ANOVA using Tukey's post hoc analysis

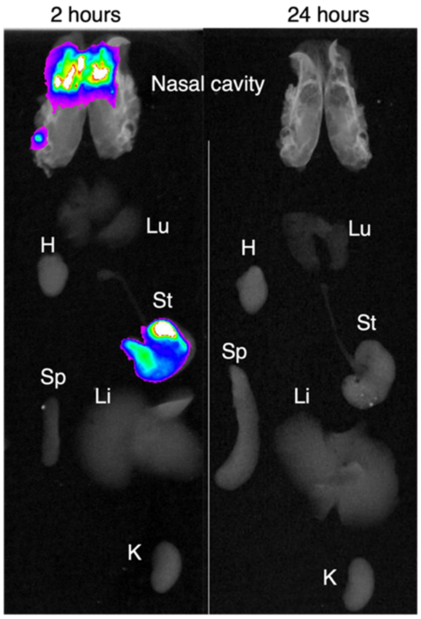


**Figure S9.** *In vivo* biodistribution of DiR-loaded PCANS. Left panel shows localization of formulation in the nasal cavity at 2 h and the right panel shows whole-body clearance of PCANS at 24 h. Lu-lungs, H-heart, St-stomach, Li-liver, Sp-spleen and K-kidney.

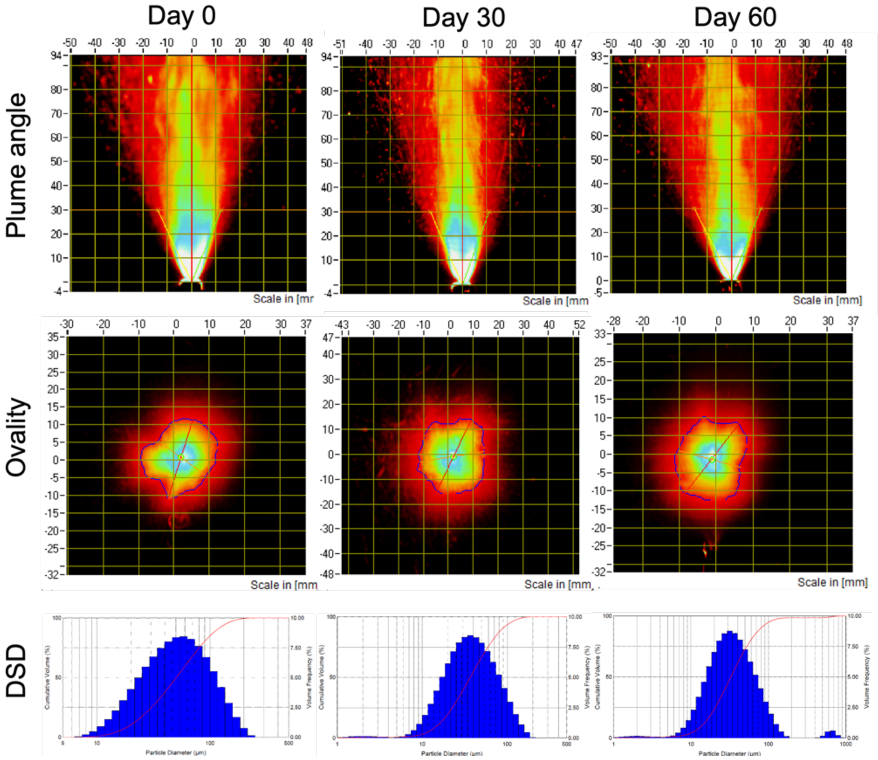


**Figure S10.** Spray characteristics of PCANS at different time points during stability study under 40°C storage conditions.

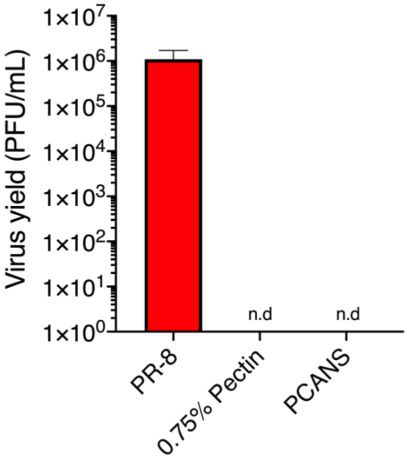


**Figure S11.** Viral load in the host cells (MDCK) after 10 min incubation of the mouse-adapted lethal influenza strain, PR-8 with pectin or PCANS.

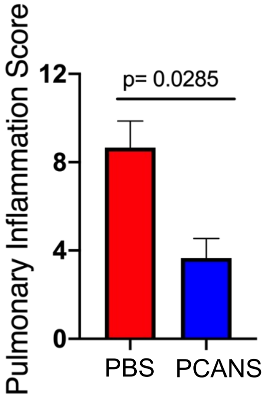


**Figure S12.** Inflammatory scoring of lung tissue of mice challenged with PR-8 and prophylactically treated with PBS or PCANS (n = 4, per group)

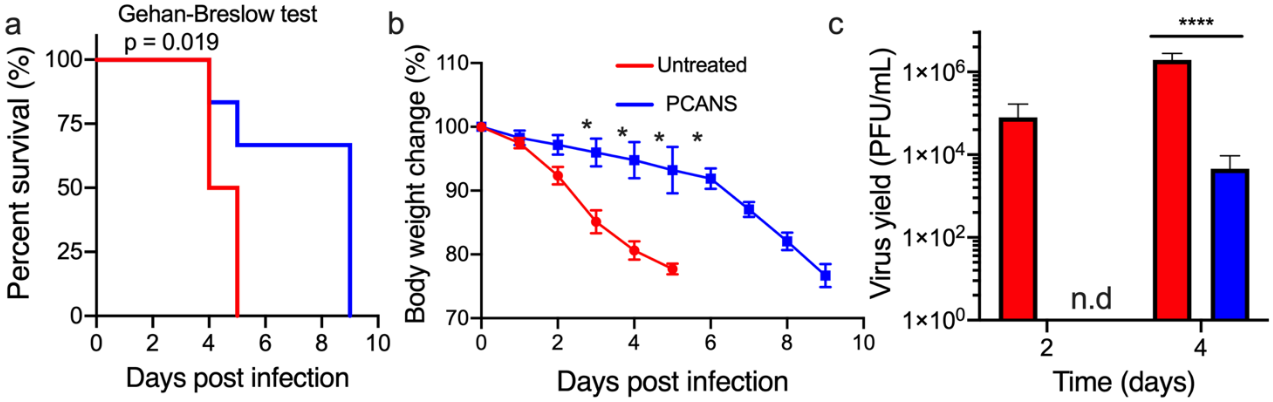

**Figure S13.** Viral titer escalation to study prophylactic efficacy of PCANS a murine model of infection. Mice received a single dose of PCANS or PBS before 15 min of intranasal infection with 500 PFU Influenza A/PR/8/34. **(a)** Survival and **(b)** body weight change of mice over a period of 10 days post-infection. *P* = 0.019 compared to the PBS-treated group for Kaplan-Meier survival curve.* *P* < 0.05. **(c)** Lung viral titer from mice on days 2 and 4 post-infection quantified by plaque assay. ****$P$<0.0001. n=6 mice/group for **a.** Data in **b** are presented as Means ± SEM (n=6 mice/group). ). Data in **c** are presented as Means ± SEM (n=4 mice/group).
